# Msh2-Msh3 interferes with DNA metabolism *in vivo*

**DOI:** 10.1101/2022.09.06.506750

**Authors:** Melisa Medina-Rivera, Samantha Phelps, Madhumita Sridharan, Jordan Becker, Natalie A. Lamb, Charanya Kumar, Mark D. Sutton, Anja Bielinsky, Lata Balakrishnan, Jennifer A. Surtees

## Abstract

Mismatch repair (MMR) is a highly conserved DNA repair pathway that safeguards the genome from errors in DNA replication. In *Saccharomyces cerevisiae*, two MutS homolog (Msh) complexes, Msh2-Msh3 or Msh2-Msh6, initiate MMR. Msh2-Msh3, the focus of this study, recognizes and directs repair of insertion/deletion loops (IDLs) up to ~17 nucleotides. Msh2-Msh3 also recognizes and binds distinct looped and branched DNA structures with varying affinities, thereby contributing to genome stability outside post-replicative MMR through homologous recombination, double-strand break repair (DSBR), and the DNA damage response. Msh2-Msh3 also promotes genome instability through trinucleotide repeat (TNR) expansions. This non-canonical activity is likely an unfortunate consequence of Msh2-Msh3’s intrinsic ability to bind a wide range of DNA structures, including those formed with single-stranded (ss) TNR sequences. We previously demonstrated that Msh2-Msh3 binding to 5’ ssDNA flap structures interfered with the *in vitro* binding and cleavage activities of the flap endonuclease Rad27 (Fen1 in mammals), which promotes 5’ ssDNA flap processing during Okazaki fragment maturation (OFM) and long-patch base excision repair (LP-BER). Here we demonstrate that elevated Msh2-Msh3 levels interfere with DNA replication and LP-BER *in vivo*, consistent with the hypothesis that protein abundance and Msh3 ATPase activities are key drivers of Msh2-Msh3-mediated genomic instability.

## INTRODUCTION

Mismatch repair (MMR) is a specialized DNA repair pathway known for its role in identifying and directing the correction of errors that evade the intrinsic fidelity mechanisms of the replication machinery, thereby increasing the fidelity of replication ~100–1000-fold (1–4). In *S. cerevisiae*, two heterodimeric MutS homolog (Msh) complexes initiate MMR with distinct but partially overlapping binding affinities (5,6). Msh2-Msh6 predominantly binds and directs the repair of single base mispairs (with the exception of C-C mismatches) and 1-2 nucleotide insertion-deletion loops (IDLs) (6–10). Msh2-Msh3 binds and directs repair of some mispairs, including A-A, C-C, and T-G (10–12), as well as both short and longer IDLs of up to 17 nucleotides in length (13–16). Following mismatch recognition, MutL homologs (Mlh), Mlh1-Mlh3, and/or Mlh1-Pms1 (Pms2 in humans) are recruited by MSH-DNA complexes in an ATP-dependent manner. The endonuclease activity of Mlh homologs is directed to cleave the nascent strand distal to the mismatch, an activity that requires a Msh complex. Mlh1-Pms1 is also activated by proliferating cell nuclear antigen (PCNA) (17–19), while Mlh1-Mlh3 is not (20,21). Mlh1-Mlh2 lacks endonuclease activity and acts as an accessory factor (22,23). Subsequently, Exo1 and replicative DNA polymerase delta (Pol δ), or epsilon (Pol ε), are recruited to remove the error and resynthesize the DNA to restore the structure of the double helix (24–26).

Msh2-Msh3 is a structure-specific DNA-binding protein that binds a variety of different DNA intermediates, with a preference for substrates with double-strand (ds)/single-strand (ss) DNA junctions (13,14,27,28). This allows Msh2-Msh3 to initiate several pathways in DNA metabolism in addition to MMR. During 3’ non-homologous tail removal (3’NHTR), a step that occurs in a sub-class of DSBR (29–32). Msh2-Msh3 binds to ds/ssDNA junctions with 3’ ssDNA non-homologous tails to stabilize it and recruits the structure-specific endonuclease Rad1-Rad10/Saw1, thereby promoting cleavage of the unannealed tails, allowing repair to proceed via DNA synthesis (29,30,33). Msh2-Msh3 also promotes heteroduplex rejection, preventing recombination between homologous sequences (34,35). In this context, Msh2-Msh3 binds IDL structures, similar to MMR, but recruits Sgs1 to unwind the D-loop. (36,37). Given these known functions, it is not surprising that loss of Msh2-Msh3 is associated with an increase in genomic instabilities that contribute to hereditary and sporadic cancers in humans (38–50). At the same time, Msh2-Msh3 also promotes genome instability in structure-specific contexts. One notable example is trinucleotide repeat (TNR) sequences; Msh2-Msh3 binding promotes the expansion of (*CNG*) tracts and likely other repeat sequences that form secondary structures (51–57), including in *S. cerevisiae* (28,58,59). Similarly, Msh2-Msh3 binding to B-DNA/Z-DNA junctions promotes mutation (60). Overexpression of Msh3 results in a base-base mismatch repair deficiency that has been attributed to an imbalance of the relative protein ratios between Msh2-Msh3 and Msh2-Msh6 (61,62).

Msh2-Msh3 ATPase activity is required to promote genome stability through MMR, 3’ NHTR, and heteroduplex rejection (63,64) and to promote genome instability through TNR expansions (65), although it is dispensable for DNA structure binding (14,64,66). Like MutS and Msh2-Msh6, Msh2-Msh3 contains two composite ATP-binding/hydrolysis sites with highly conserved Walker A and Walker B adenosine nucleotide-binding sites that are essential for ATP binding and hydrolysis, respectively (67–70). In *S. cerevisiae*, amino acid substitutions of the Walker A (G796 in Msh3), predicted to prevent ATP binding, abolished Msh2-Msh3-mediated MMR (64). Notably, Msh2-Msh3 binding to different DNA structures alters the kinetics of Msh2-Msh3 ATP binding, hydrolysis, and nucleotide turnover, which impact downstream steps such as Msh2-Msh3 turnover and recruitment of partner proteins, promoting genome stability or instability (14,27,28,58,66,68,71,72).

5’ ssDNA flaps are generated in at least two DNA metabolic pathways: Okazaki fragment maturation (OFM) during DNA replication and long-patch base excision repair (LP-BER) (**Figure 1**). Polymerase (Pol) δ extends the initiator primer in OFM and primer upstream of the abasic site in LP-BER, eventually encountering the preceding DNA fragment. Pol δ proceeds with synthesis, displacing the 5’-end of the downstream segment, forming a single-stranded 5’ flap structure (73–75). This intermediate is cleaved by endonuclease Rad27 (Fen1 in mammals), leaving a nick that is sealed by DNA ligase Cdc9^LigI^ (DNA Ligase I (LigI) in humans). We previously demonstrated that Msh2-Msh3 binds 5’ ssDNA flap structures, albeit with lower affinity than 3’ ssDNA flaps (14), to form a specific complex that interacts with the ss/dsDNA junction (28). We demonstrated that Msh2-Msh3 competes with both Rad27^FEN1^ and Cdc9^LigI^ for binding to DNA substrates. This resulted in the inhibition of Rad27 ^FEN1^ endonuclease activity, ligation, and a significant reduction of Okazaki fragment processing *in vitro* (28). Given that 5’ flap processing is essential for DNA metabolism *in vivo*, uncontrolled binding of Msh2-Msh3 to 5’ flap intermediates poses a potential risk for normal DNA metabolism.

**Figure 1.**
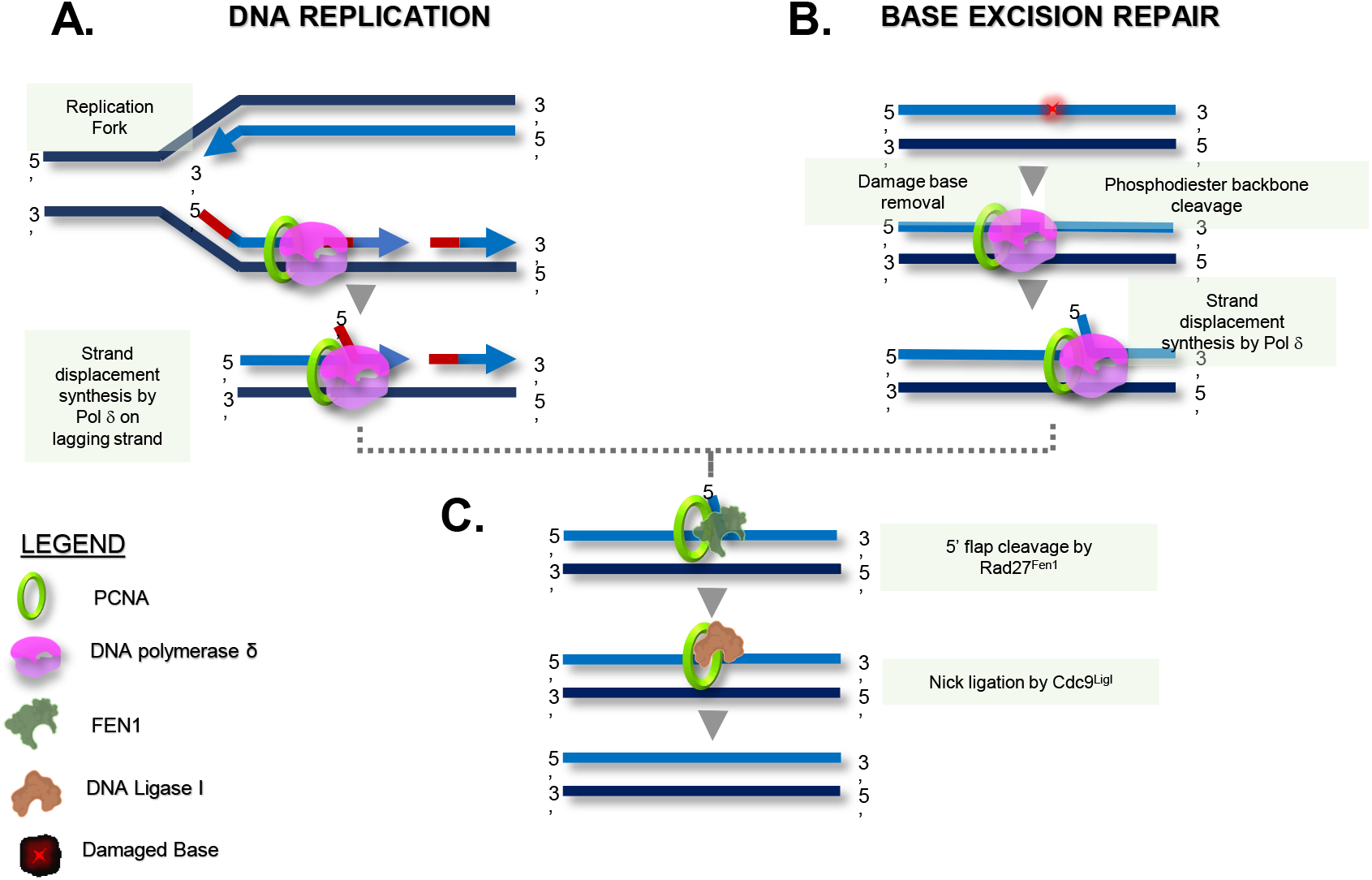
Two DNA metabolic pathways that involve 5’ ssDNA flap processing. **(A)** During lagging strand synthesis DNA polymerase δ (Pol δ) initiates strand displacement synthesis when it encounters the upstream Okazaki fragment. **(B)** Base excision repair (BER) pathway is the primary mechanism to remove alkylation (and other) DNA damage. In long patch BER, DNA glycosylase catalyzes cleavage of the N-glycosidic bond of a damaged base to generate an apurinic/apyrimidinic (AP) site. AP endonuclease cleaves the AP site to generate a gap intermediate in which two or more (<10) nucleotides are removed. If polymerase d is recruited for gap filling synthesis, this will lead to strand displacement. **(C)** In both pathways, a 5’ ssDNA flap structure is formed. Rad27 binds and cleaves the flap and Cdc9 ligates the resulting nick, leading to either a continuous lagging strand or repaired DNA.

Here we present evidence that elevated levels of Msh2-Msh3 interfere with DNA metabolism *in vivo*. Even low levels of Msh2-Msh3 overexpression increased sensitivity to the alkylating drug methyl methanesulfonate (MMS), which generates lesions typically repaired by base excision repair (BER). Msh2-Msh3 overexpression also induced defects in cell cycle progression that are likely a result of Okazaki fragment stress. Our results support a model in which elevated levels of Msh2-Msh3 interfere with normal DNA metabolism, not simply by binding DNA substrates but by engaging in aberrant signaling in an ATP binding-dependent manner. Furthermore, they suggest that tight regulation of the Msh2-Msh3 expression levels is important *in vivo* to prevent interference with both DNA synthesis and the processing of a variety of DNA structures.

## MATERIALS AND METHODS

### Plasmids and Yeast Strains

To generate low-copy and high-copy plasmids expressing *MSH3,* a *Sac*II-*Pst*I fragment from pEAI215 (71), which includes DNA sequence from ~ 1 kb upstream of *MSH3* to ~ 500 bp downstream of *MSH3* from its endogenous chromosomal location, was ligated into pRS423 (76) digested with *Sac*II and *Pst*I to generate pSP1. This is a high copy 2μ plasmid with a *LEU2* marker. From this plasmid, a *Sac*II-*Sal*I fragment was excised, containing the entire *MSH3* sequence from pEAI215, and ligated into pRS424 (76) and pRS414 (77) digested with *Sac*II and *Sal*I to generate pSP15 and pSP18, respectively. Both carry a *TRP1* marker; pSP15 is a 2μ plasmid, while pSP18 is a single copy *ARS CEN* plasmid. Plasmids were transformed into a *msh3*Δ yeast strain background using the lithium acetate method (78).

Overexpression plasmids of *MSH2* (pMMR8) and *msh2G693D* (pEAE270) (**Table S1**) have been described previously (13,14,79). Galactose-inducible overexpression plasmids of *MSH3* (pMMR20), *msh3Y925A* (pCK94 or pMME2), *msh3G796A* (pCK42), *MSH6* (pEAE218), and empty vector (pJAS104) were described previously (**Table S1**) (13,14,33,64,79). We generated *msh3D870A* in pEAI218 (71) by site-directed mutagenesis. A *Bsu36I*-*MluI* fragment from this plasmid, containing the *msh3D870A*, was sub-cloned into pMMR20 (13), to generate a galactose-inducible *msh3D870A* overexpression plasmid (pMME3). *MSH3*, *msh3,* or empty vector plasmids (all carrying the *leu2D* nutritional marker) were co-transformed with the *MSH2* or *msh2* overexpression plasmid (*TRP1* marker) into various yeast strains using the lithium acetate method (78). For the His-PCNA (*POL30*) and His-pcna (*pol30*) mutant experiments, yb2062 (*His-POL30*), yb2063 (*His-pol30K164R*), yb2064 (*His-pol30K242R*), yb2066 (*His-pol30K164R/K242R)*(80) were made *trp1-* and *leu2-* by sequential marker swap by *hisG-URA3-hisG* pop-out with pNKY85 (targeting *LEU2*) and pNKY1009 (targeting *TRP1*) (81,82) to generate JSY4937-4945 (*His-POL30*), JSY5007-5012 (*His-pol30K164R*). JSY5013-14 (*His-pol30K242R*) and JSY5015-27 (*His-pol30K164R/K242R*). These strains were co-transformed with pMMR8 and pMMR20 or pJAS104.

All strains used in this study are described in **Table S2**.

### Galactose Inducible Overexpression

Cultures of a *msh3Δ* (JSY1505 or JSY905) or *His-POL30*/*pol30* backgrounds carrying both pMMR8 (*MSH2*) or pMMR8-derived (*msh2G693D*) and pMMR20 (*MSH3*) or pMMR20-derived (*msh3* alleles) (13) were grown to mid-log phase in synthetic complete (SC) medium in the presence of 2% lactate and 2% glycerol as carbon sources. Protein expression was induced by the addition of 2% galactose for 17 hours. Uninduced and induced cells were collected for flow cytometry, quantitative real-time PCR (qRT-PCR), and/or western blotting to analyze PCNA modification. Cells harvested for flow cytometry were washed with sterile deionized water and fixed in 70% ethanol at 4°C for a minimum of 1 hour (up to 1 week) before flow cytometry analysis. Cells harvested for RNA extraction were washed with UltraPure™ DNase/RNase-Free Distilled Water (Invitrogen), harvested by centrifugation, and resuspended in β-mercaptoethanol/Buffer RLT solution as described by RNeasy Mini Kit (QIAGEN) guidelines. Aliquots from each time point were snap-frozen in liquid nitrogen and stored at −80°C until ready for processing.

### RNA Isolation and Quantitative Real-Time PCR

Total RNA was isolated from cultured yeast cells using the QIAGEN RNeasy Mini Kit. As recommended by the manufacturer, residual DNA was removed by on-column DNase I digestion was carried out for 15 minutes. One μg of total RNA was reverse transcribed using a mix of oligo(dT) and random hexamer primers following the manufacturer’s instructions of the iScript^TM^ cDNA synthesis kit (BioRad). Primers to detect endogenous transcript levels of *MSH2*, *MSH3*, *MSH6,* and *PDA1* are described in **Table S3**. The RT-PCR was performed at 95 °C for 3 minutes, forty cycles of amplification consisting of denaturation at 95 °C for 15 seconds, annealing at 55 °C for 30 seconds, extension at 72 °C for 30 seconds, followed by melting curve analysis. Each RNA extraction was performed a minimum of three times. Each PCR included a standard curve with genomic DNA, and the levels of target transcripts were normalized to that of the reference gene that encodes Pyruvate Dehydrogenase Alpha 1 (*PDA1*).

### Canavanine Resistance Assays

Mutation rates were measured at the *CAN1* locus as previously described (83,84). Briefly, strains were grown on SC–Trp–Leu plates until colonies reached 2 mm in size. The carbon source in the plates was 2% glucose or 2% galactose for *MSH3* uninduced or induced, respectively. Colonies were then suspended in 100 μL of 1x TE (10 mM Tris-HCl, pH 7.4; 1 mM EDTA) and diluted 1:10,000. Twenty μL of the undiluted colony suspension was plated on SC–Trp–Leu–Arg + Canavanine, and 100 μL of the 10^−4^ dilution was plated on SC–Trp–Leu–Arg. Both permissive and selective plates contained 2% glucose as a carbon source. The plates were incubated at 30°C until colonies reached ~1-2 mm in size. Colonies were counted, and mutation rates and 95% confidence intervals were calculated through FluCalc fluctuation analysis software (85). Assays were performed on multiple independent isolates for each genotype on separate days.

### Methyl Methanesulfonate Survival Assays

Cultures were grown to the mid-log phase in liquid SC–Trp media for assays performed with high and low-copy plasmids. Cells were diluted and plated on appropriate SC–Trp plates or SC–Trp plates containing 0.005, 0.01, 0.015, 0.0175, or 0.020% MMS. To avoid degradation of MMS and ensure consistent results, cells were plated within hours of pouring the plates. After incubation at 30°C for 4 days, percent survival was calculated as the ratio of the number of colonies that grew in the presence of MMS relative to the no MMS control. Assays were repeated a minimum of three times with at least two independent isolates.

For assays performed with overexpression strains, cultures were grown to the mid-log phase in SC media in the presence of lactate and glycerol as carbon sources. As described above, *MSH3* or *msh3* expression was induced by adding 2% galactose. Cells were diluted and plated into appropriate SC–Trp–Leu plates or SC–Trp–Leu plates containing 0.005% MMS.

### Cell Cycle Analysis by Flow Cytometry

After fixation in 70% ethanol, yeast cells were washed with sodium citrate/EDTA solution (50 mM sodium citrate and 1 mM EDTA [pH 8.0]) and treated with 0.06 mg RNase A (Invitrogen) at 50°C for 2 hours. This was followed by adding 0.25 mg of Proteinase K (Sigma-Aldrich) and incubation at 50°C for 1-2 hours. Cells were mixed with a sodium citrate/EDTA solution containing 1 μM SYTOX® Green. Stained DNA was then analyzed for chromosomal content using a BD Fortessa Flow Cytometer at an excitation wavelength of 488 nm. Data shown were analyzed using BD FACSDiva^TM^ and FlowJo^TM^ software.

### Detection of PCNA Modification by Western Blot

Total protein extracts from yeast strains overexpressing Msh2-Msh3 were TCA precipitated and analyzed by Western blot with an anti-PCNA antibody as described previously (86,87). Blots were imaged with Bio-Rad Chemi-Doc Touch Imaging System. Linear changes in exposure were applied to entire blots. PCNA signal intensity was quantified using ImageJ software.

### DNA Substrates

Oligonucleotides were synthesized by Integrated DNA Technologies (IDT, Coralville, IA) or Midland Certified Reagents Company (TX). For ATPase assays, the homoduplex (LS1/LS2) and +8 loop MMR (LS2/LS8) substrates sequences, assembly and purification were as described previously (14,66,71). Radiolabeled isotope was purchased from Perkin Elmer Life Sciences. Synthetic oligonucleotides were labeled with [γ ^32^P]-ATP and T4 polynucleotide kinase (New England Biolabs) at their 5’ end, as previously described (88). Following gel purification, substrate annealing was performed in a 1:2:4 ratio (labeled oligonucleotide: template: second oligonucleotide), as previously described (88). The oligonucleotides used in this study are listed in **Table S4**.

### Protein purification

*S. cerevisiae* Msh2-Msh3 (66) and Pol δ (89) were expressed and purified as previously described.

### Pol δ Strand Extension Assay

Two DNA substrates were used in the strand extension assays: the synthesis substrate and the strand displacement substrate. The synthesis substrate was created by annealing 5’ ^32^P-labeled 44mer to 110nt template sequence in a 1:2 ratio, respectively. The strand displacement substrate was formed by annealing a 5’ ^32^P-labeled 44 nucleotides upstream primer to a 110nt template containing a 60nt downstream primer in a 1:2:4 ratio. Five nM of the synthesis substrate was incubated with Pol δ (75nM with the synthesis substrate and 150 nM for the strand displacement substrate) and increasing concentrations of Msh2-Msh3 (50 nM, 100 nM, 250 nM) for 30 minutes at 30°C. Reactions were performed in 20 µL volume in reaction buffer containing 50 mM Tris-HCl pH 8.0; 2 mM DTT, 2 µg/mL bovine serum albumin, 8 mM MgCl_2_, 1 mM ATP, 0.1 mM dNTPs, 25 nM NaCl. Reactions were terminated using 2X termination dye containing 90% formamide (v/v), 10 mM EDTA, with 0.01% bromophenol blue and xylene cyanole. Samples were then resolved on a 12% polyacrylamide gel containing 7M urea. Products were analyzed by a PhosphorImager (Typhoon 9500) and quantified using ImageQuant version 5.2 (Molecular Dynamics). All experiments were done at least in triplicate. Representative gels are shown.

### ATPase Assays

Hydrolysis of ATP was monitored using a coupled spectrophotometric assay as described previously (66,90). In this assay, the conversion of ATP to ADP and *P_i_* is linked to the oxidation of NADH to NAD^+^ and is monitored as a decrease in absorbance at 340 nm. Assays were performed at 30°C and monitored using a Varian, Cary-50 Bio UV–vis spectrophotometer. The reactions contained 20 mM Tris-acetate (pH 7.5), 0.3 mM NADH, 5 mM PEP, 20 U/mL pyruvate kinase, 20 U/mL lactate dehydrogenase, 2 mM magnesium acetate, DNA (250 nM) and Msh2–Msh3 (50 nM) and up to 2.5 mM ATP. Msh2-Msh3 was pre-bound to DNA, followed by adding ATP in small increments. Approximately 80 data points were fit to a linear curve. The rate of ATP hydrolysis at each ATP concentration was calculated by multiplying the slope of the line by 159 (the change in absorbance of NADH per unit time) (90).

## RESULTS

### Characterization of *MSH3* expression levels

Msh2-Msh3 binding interferes with the processing of 5’ ssDNA flap intermediates by Rad27^FEN1^ and Cdc9^LigI^ *in vitro*, competing with Rad27^FEN1^ for binding the DNA intermediate (28). Because 5’ ssDNA flaps are generated in multiple DNA metabolic pathways, such as lagging strand synthesis, and long-patch BER, we were intrigued by the possibility that Msh2-Msh3 might interfere in these processes *in vivo,* leading to genome instability and replication stress. Notably, elevated levels of Msh2-Msh3 are a key driver of genome instability in eukaryotes *via* TNR expansions (91,92). Based on these results, we set out to test the hypothesis that elevated Msh2-Msh3 levels can drive genomic instability *via* 5’ flap DNA intermediates.

To address this hypothesis, we established a series of plasmids to express *MSH3* at different levels. First, we generated a single copy *ARS CEN* plasmid carrying *MSH3* under the control of its endogenous promoter (pSP18, *MSH3-LC*). Second, we constructed a high copy 2μ plasmid carrying *MSH3* under the control of its endogenous promoter (pSP15, *MSH3-HC*). Finally, *MSH2* and *MSH3* were co-overexpressed as described previously; *MSH2* (pMMR8) expression was under the control of the *ADC1* promoter, expressed constitutively and at elevated levels, and *MSH3* (pMMR20) was expressed under the control of a galactose-inducible *GAL-PGK* promoter (13,14).

The endogenous Msh3 protein levels are not detectable by Western blot (64). Instead, we measured expression levels by qRT-PCR, although we note that there is not necessarily a linear relationship between gene expression and protein levels. RNA was extracted from wild-type and *msh3*Δ cells. Consistent with low endogenous Msh3 protein levels, we observed endogenous *MSH3* mRNA levels to be six times lower than endogenous *MSH2* or *MSH6* expression levels (**Figure 2A,B**). The *MSH3* mRNA levels were four and sixty-four times higher than *MSH3* endogenous levels when expressed from *MSH3-LC* or *MSH3-HC* plasmids, respectively (**Figure 2A,B**). No significant differences in *MSH2* or *MSH6* expression levels were observed in the presence of *MSH3-LC* or *MSH3-HC*. (**Figure 2B**).

**Figure 2.**
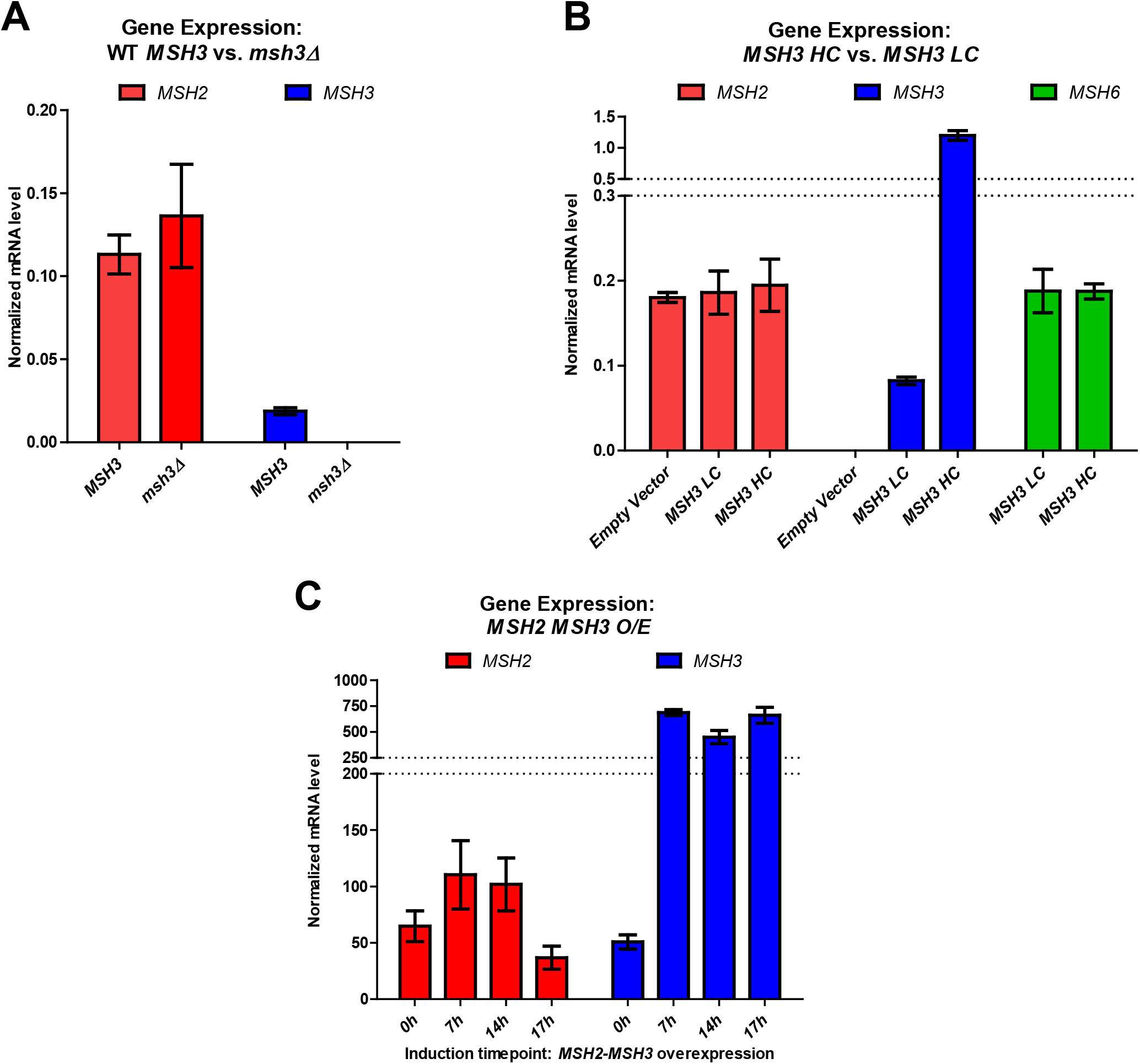
Expression levels of *MSH2*, *MSH3* and *MSH6* in different cellular contexts. Endogenous mRNA levels of *MSH2* (red), *MSH3* (blue), and *MSH6* (green) were measured using RT-qPCR. **(A)** RNA was isolated from *MSH3* or *msh3*Δ yeast cells at mid-log phase. **(B)** RNA was isolated at mid-log phase from *msh3*Δ strains carrying either an empty vector or either a low copy number (*ARS CEN;* LC) or a high copy number (*2 m;* HC) plasmid, bearing *MSH3* under the control of the endogenous *MSH3* promoter. **(C)** RNA was isolated from strains co-overexpressing *MSH2,* under a constitutive promoter, and *MSH3,* under a galactose-inducible promoter. Culture aliquots were collected at indicated hours after induction with galactose. All RNA levels were normalized to the reference gene *PDA1.* Data represents the mean of at least three independent experiments. Error bars represent SEM.

The *msh3*Δ strain was co-transformed with *MSH2* (pMMR8), and a second plasmid bearing either a galactose-inducible copy of *MSH3* (pMMR20) or an empty vector control derived from pMMR20 (pJAS104 [EV]). Prior to galactose induction, we observed an increase in *MSH3* mRNA of approximately ~2,300-fold over *MSH3* endogenous levels in these strains (**Figure 2A,C**), indicating read-through transcription in the absence of galactose. No *MSH3* mRNA was observed in the control *msh3*Δ strain co-expressing *MSH2* and the empty vector (pJAS104) (**Figure S1**). After induction by galactose for 17h, *MSH3* expression levels increased to ~14-fold compared to pre-induction (0h) levels, ~32,000 times higher than the endogenous levels of *MSH3* mRNA (**Figure 2A,C**). Using these constructs, we previously demonstrated detectable Msh3 or msh3 protein levels following induction, including *msh3* alleles tested here, although protein levels are still low (64).

Studies in human cell lines demonstrated a strong Msh2-Msh6-specific mutator phenotype when *MSH3* is overexpressed, presumably because excess Msh3 competes with Msh6 for interactions with Msh2 (61,62). To functionally test *MSH3* overexpression in yeast, we performed canavanine resistance assays, a Msh2-Msh6-specific mutator assay following galactose-induced overexpression of *MSH3* (**Table 1**). *MSH3* overexpression (*O/E*) in galactose increased the mutation rate ~9-fold over the rate observed in *MSH3 O/E* grown in glucose, similar to the elevated mutation rate in *msh6*Δ, consistent with the presence of elevated Msh3 protein. Growth of *WT* strains in galactose in the absence of *MSH3 O/E* did not have this effect (**Table 1**). *MSH3-HC* also increased the mutation rate, albeit to a lesser extent (~2-fold), while *MSH3-LC* had no effect (**Table 1**).

**Table 1.**
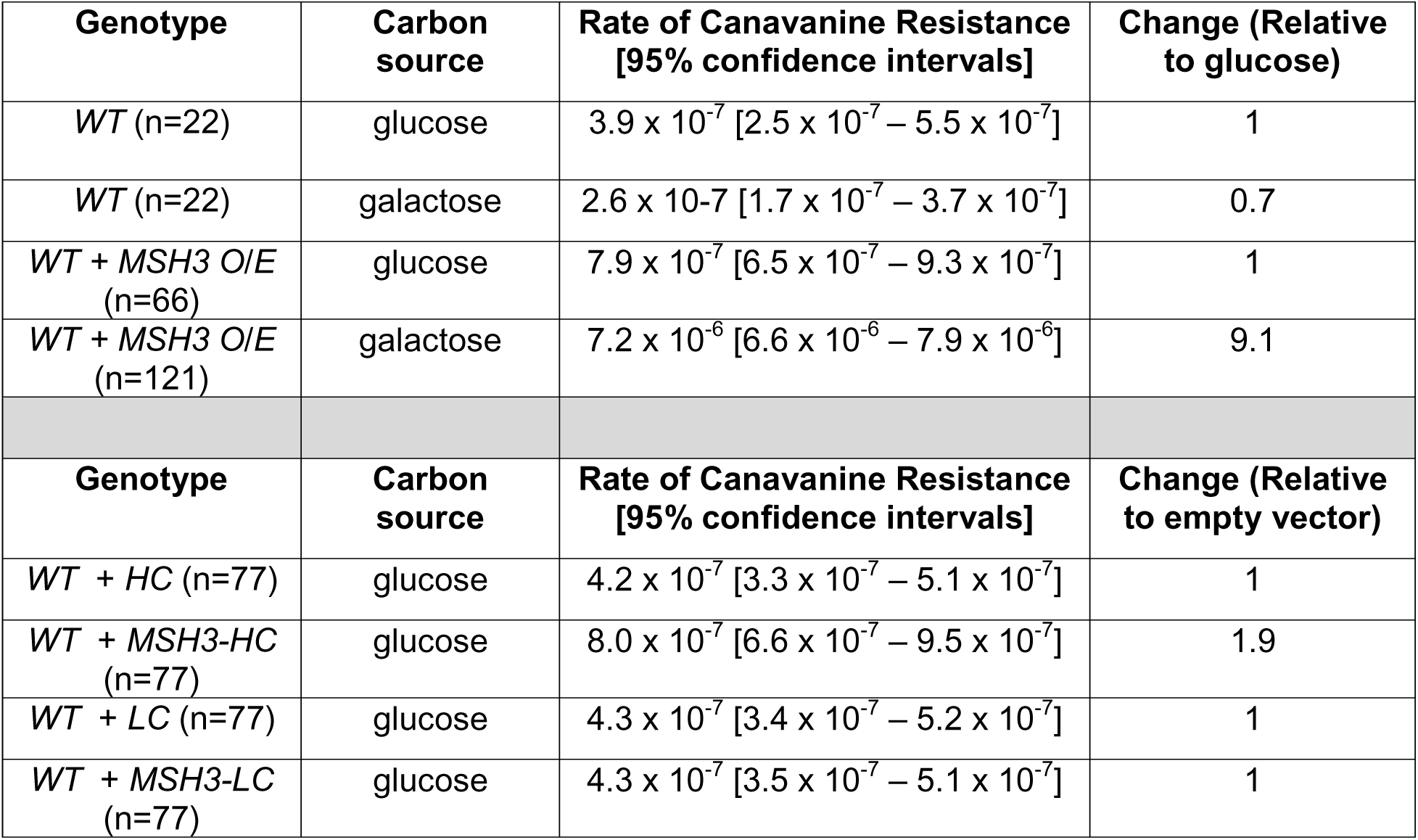
Canavanine resistance mutation rates.

| Genotype | Carbon source | Rate of Canavanine Resistance [95% confidence intervals] | Change (Relative to glucose) |
| --- | --- | --- | --- |
| *WT* (n=22) | glucose | $3.9 \times 10^{-7}$ [$2.5 \times 10^{-7} - 5.5 \times 10^{-7}$] | 1 |
| *WT* (n=22) | galactose | $2.6 \times 10^{-7}$ [$1.7 \times 10^{-7} - 3.7 \times 10^{-7}$] | 0.7 |
| *WT + MSH3 O/E* (n=66) | glucose | $7.9 \times 10^{-7}$ [$6.5 \times 10^{-7} - 9.3 \times 10^{-7}$] | 1 |
| *WT + MSH3 O/E* (n=121) | galactose | $7.2 \times 10^{-6}$ [$6.6 \times 10^{-6} - 7.9 \times 10^{-6}$] | 9.1 |
| Genotype | Carbon source | Rate of Canavanine Resistance [95% confidence intervals] | Change (Relative to empty vector) |
| *WT + HC* (n=77) | glucose | $4.2 \times 10^{-7}$ [$3.3 \times 10^{-7} - 5.1 \times 10^{-7}$] | 1 |
| *WT + MSH3-HC* (n=77) | glucose | $8.0 \times 10^{-7}$ [$6.6 \times 10^{-7} - 9.5 \times 10^{-7}$] | 1.9 |
| *WT + LC* (n=77) | glucose | $4.3 \times 10^{-7}$ [$3.4 \times 10^{-7} - 5.2 \times 10^{-7}$] | 1 |
| *WT + MSH3-LC* (n=77) | glucose | $4.3 \times 10^{-7}$ [$3.5 \times 10^{-7} - 5.1 \times 10^{-7}$] | 1 |

### Overexpression of Msh2-Msh3 leads to MMS sensitivity

Given that Msh2-Msh3 interferes with 5’ flap processing *in vitro*, we tested whether elevated *MSH3* expression sensitized the cells to MMS, a monofunctional alkylation agent that methylates DNA at N^7^-deoxyguanine and N^3^-deoxyadenine (93), a lesion typically repaired by the BER pathway. *msh3*Δ yeast cells carrying *MSH3-LC* or *MSH3-HC* plasmids were grown to mid-log phase, serially diluted, and grown on selective (SC-trp) plates in the absence or presence of MMS. At 0.015%, 0.0175%, and 0.02% concentrations of MMS, cells carrying either *MSH3-LC* or *MSH3-HC* exhibited mild but significant sensitivity to MMS compared to empty vector controls (**Figure 3A**). When *MSH2* and *MSH3* were co-overexpressed, yeast cells exhibited significant sensitivity to 0.005% MMS compared to overexpression of *MSH2* alone (**Figure 3B**). With higher *MSH3* overexpression, cells were sensitive to much lower MMS concentrations, consistent with a correlation between Msh3 levels and MMS sensitivity. These results indicate that excess Msh2-Msh3 compromises LP-BER, suggesting interference with 5’ flap processing *in vivo*.

**Figure 3.**
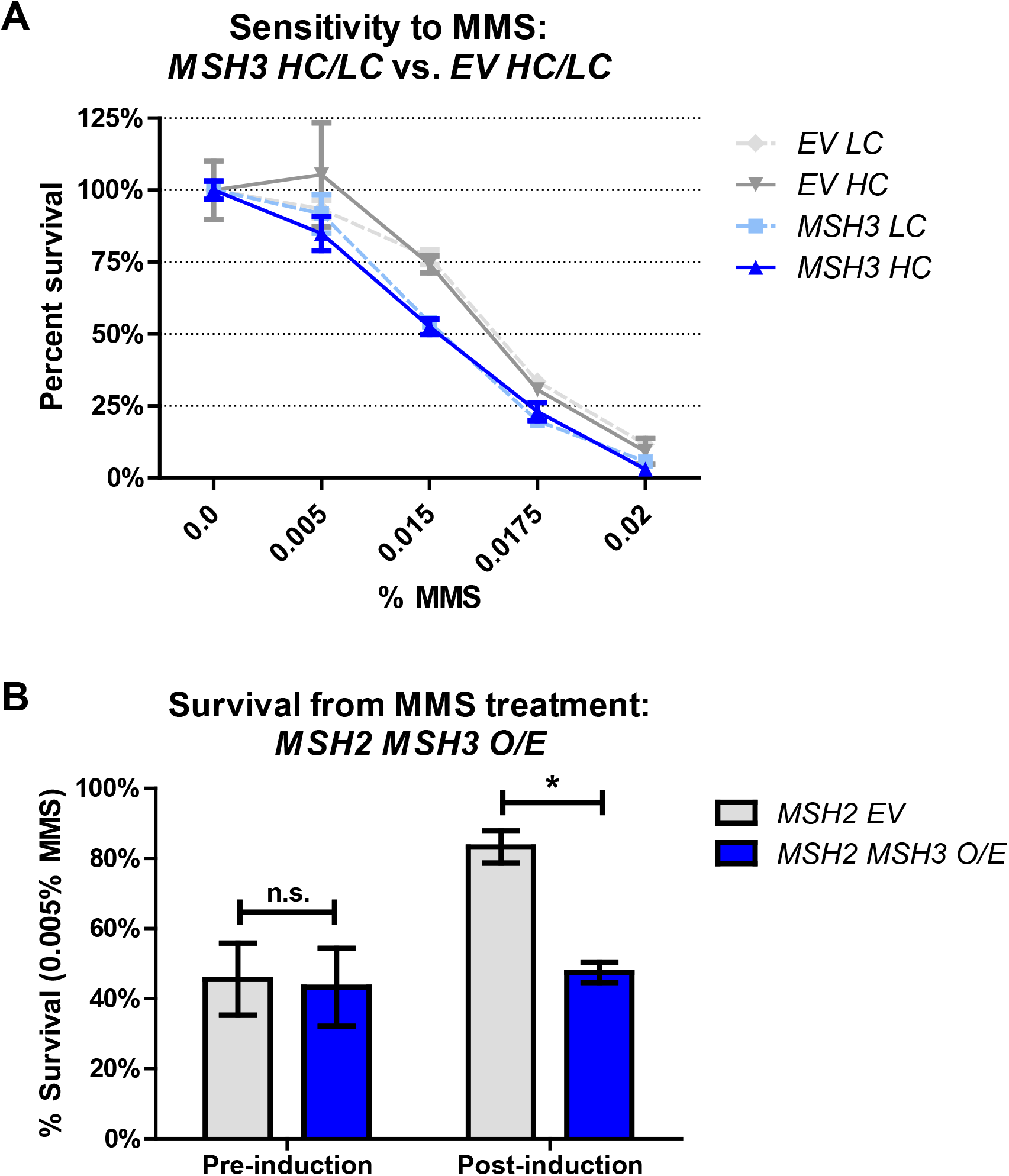
Elevated MSH3 expression renders cells sensitive to methyl methanesulfonate (MMS). **(A)** MMS sensitivity of *msh3*Δ strains carrying either a low-copy (*MSH3 LC*, light shade) or high-copy number (*MSH3 HC*, dark shade) plasmid containing either empty vector (*EV*, gray lines) or *MSH3* (blue lines) were grown in increasing concentrations of MMS. Plotted graphs represent at least two independent experiments done in triplicate and with at least two independent transformants. Error bars were calculated by SEM; p values were calculated in Prism (*MSH3 LC* vs. *EV LC* p <0.0001; *MSH3 HC* vs. *EV HC*, p = 0.0317). **(B)** MMS sensitivity of *msh3*Δ strains co-overexpressing *MSH2* and *MSH3*. Cultures were grown to mid-log phase and *MSH3* expression was induced by the addition of galactose. Time point 17 hours after induction was collected, serial diluted, and plated onto SC agar plates in the absence or presence of 0.005% MMS. Plotted data represents the results of at least three independent experiments. Error bars represent the SEM; p values were calculated in Prism (*p = 0.0297).

### Overexpression of Msh2-Msh3 interferes with cell cycle progression

Okazaki fragment maturation (OFM) also requires the processing of a displaced 5’ ssDNA flap; interference with OFM causes delays in cell cycle progression (80,87,94–97). Therefore, we assessed cell cycle progression by flow cytometry in cells overexpressing *MSH2* and *MSH3*. After *MSH2* and *MSH3* co-overexpression, we observed substantial defects in cell cycle progression, which were not observed in the presence of the empty vector or *MSH2* and *MSH6* co-overexpression (**Figure 4**). The cell cycle profile shifted dramatically, with an apparent accumulation of cells in the early S phase and a complete loss of defined G1 (1C) and G2/M (2C) populations. We quantified cell populations in G1, S, or G2/M phases of the Cell Cycle function in FlowJo software (see **Figure S2**). Upon *MSH2 MSH3* co-overexpression, we observed a substantial increase in the number of cells in the S phase with a concomitant decrease in cells in G1 and, to a lesser extent, in G2/M (**Figure 4, Figure S3**). This accumulation of cells in the S phase is typically associated with slowed fork progression and/or DNA damage (98–100). These data suggest that overexpression of Msh2-Msh3 interferes with normal DNA metabolism, slowing down or inhibiting S phase. Notably, when the cells were released back into glucose, reversing the *MSH3* induction, they eventually resumed normal cell cycle progression (**Figure 5**). Notably, the cell cycle profile of cells co-overexpressing *MSH2 MSH3* resembled that of *rad27Δ* cells at elevated temperature (97), consistent with a model in which excess Msh2-Msh3 outcompetes Rad27^FEN1^ *in vivo* and *MSH2 MSH3* overexpression makes the cells functionally *RAD27* null.

**Figure 4.**
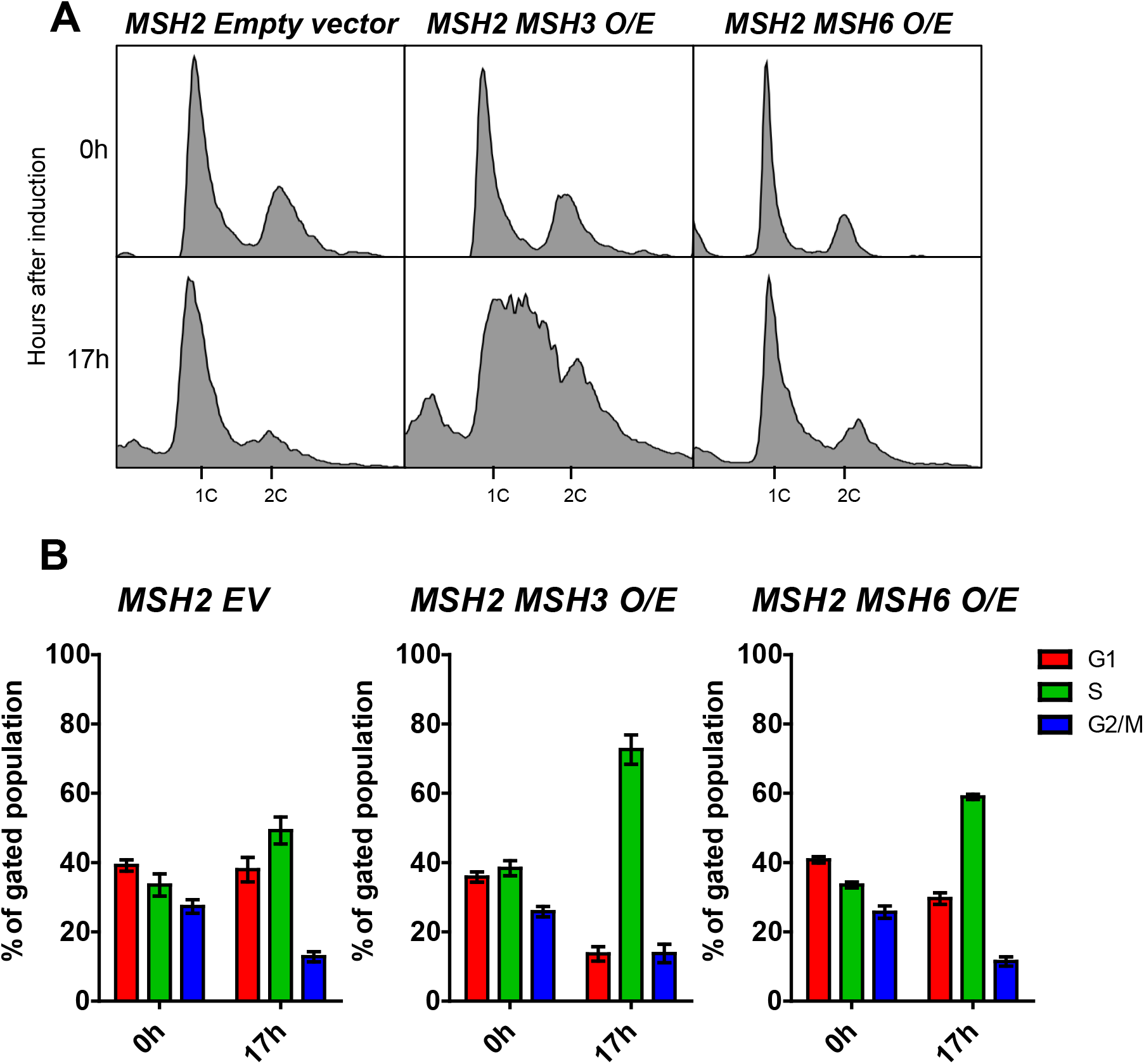
*MSH2-MSH3* overexpression induces a delay in cell cycle progression. *MSH2* and *MSH3* were overexpressed following galactose induction in a *msh3*Δ background. Aliquots were collected at 0 and 17 hours after induction. **(A)** Histograms are shown of chromosomal content of asynchronous populations of *MSH2 + empty vector (EV), MSH2 + MSH3,* and *MSH2 + MSH6* at several timepoints following addition of galactose. 1C indicates 1x DNA content; 2C indicates 2x DNA content. **(B)** Quantification of relative proportion of cells in different phases of the cell cycle for *MSH2 + EV*, *MSH2 + MSH3,* or *MSH2 + MSH6.* The percentage of cells in G1 (1C), G2/M (2C) or S (between 1C and 2C) phases was determined using FlowJo software (see Figure S1 for details). Plotted values correspond to data collected from at least three independent experiments from at least two independent isolates. Error bars represent SEM.

**Figure 5.**
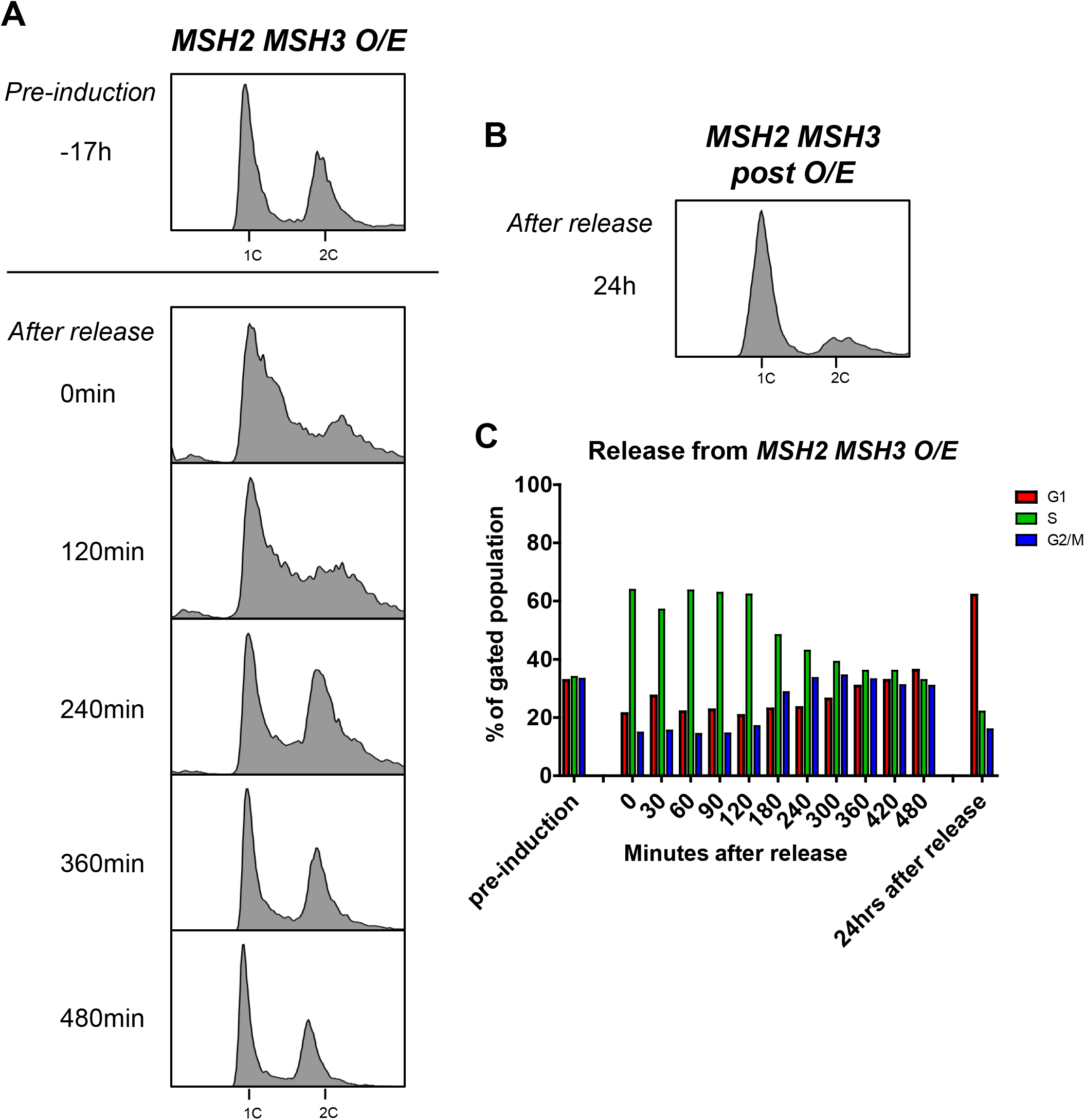
Recovery from *MSH2 MSH3* overexpression. *MSH2* and *MSH3* were overexpressed and induced as previously described. Cells were released from induction by transfer into glucose-containing media. Samples were collected and fixed for flow cytometry analysis **(A)** before induction (−17h), after induction (0 min), after release into glucose every 30-60 minutes for 8 hours, and **(B)** 24 hours after release. Histograms show the distribution of chromosomal content at each time point. **(C)** Cell cycle profiles were quantified using FlowJo software (See Figure S2).

### *MSH3* overexpression enhances PCNA post-translational modification

Interference with Okazaki fragment processing leads to cell cycle delays and DNA damage responses, signaling cascades that are partly mediated by post-translational modifications in PCNA (86,87,97). We examined whether *MSH2 MSH3* overexpression triggered PCNA modifications, which would support the hypothesis that a DNA damage response is activated. Following *MSH2 MSH3* overexpression, whole-protein TCA precipitation was performed, and the resulting cell extracts were analyzed by western blot with α-PCNA antibody (86,87). We observed two PCNA-specific bands, one consistent with the size of unmodified PCNA (~29 kDa) (**Figure 6A**) and a PCNA band with lower mobility (~49 kDa), consistent with a post-translationally modified form of the protein. This modified PCNA band was significantly increased following *MSH2 MSH3* overexpression compared to the empty vector (**Figure 6B**) and *MSH2 MSH6* overexpression (**Figure S4**).

**Figure 6.**
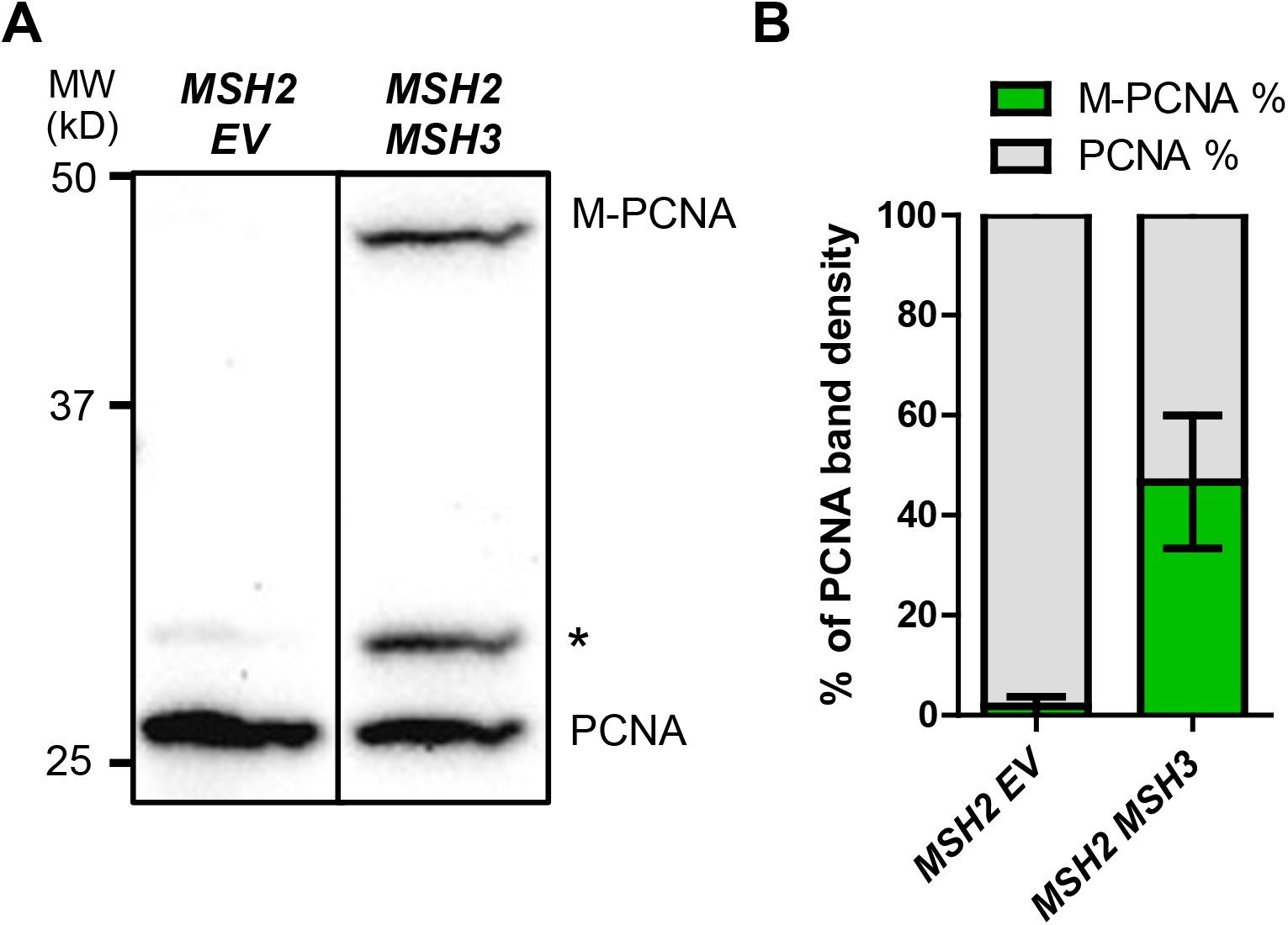
*MSH2 MSH3* overexpression induces post-translational modification of PCNA. *MSH2* and *MSH3* or empty vector were overexpressed in a *msh3*Δ background, as previously described. Following induction, TCA protein extracts were prepared, and the proteins were separated by 12% SDS-PAGE, transferred onto a membrane and then probed with anti-PCNA (P4D1). **(A)** Western Blot for PCNA. Modified PCNA is marked as M-PCNA. The asterisk indicates non-specific bands. These data are from a single blot, but not adjacent lanes, as indicated by the vertical line. **(B)** PCNA Western Blot quantification, shown as relative band densities of modified versus unmodified PCNA in terms of the percentage of total PCNA signal (measured as the sum of unmodified and modified bands).

PCNA is differentially modified in response to distinct signaling pathways (86,101,102). To determine the residue at which PCNA is modified following *MSH2 MSH3* overexpression, we compared the modification of His-PCNA, His-pcnaK164R, His-pcnaK242R, and His-pcnaK164R/K242R. Mutation of these highly conserved lysines to arginine renders them unmodifiable with either SUMO or ubiquitin. When *MSH2 MSH3* was overexpressed in the presence of *His-pol30-K164R* or *His-pol30-K164/RK242R*, modified PCNA was no longer detectable. The *His-pol30-K242R* exhibited modification levels similar to His-PCNA (**Figure 7C**). These results indicated that the *MSH2 MSH3* overexpression-dependent post-translational modification of PCNA occurs at K164 PCNA and not at K242.

**Figure 7.**
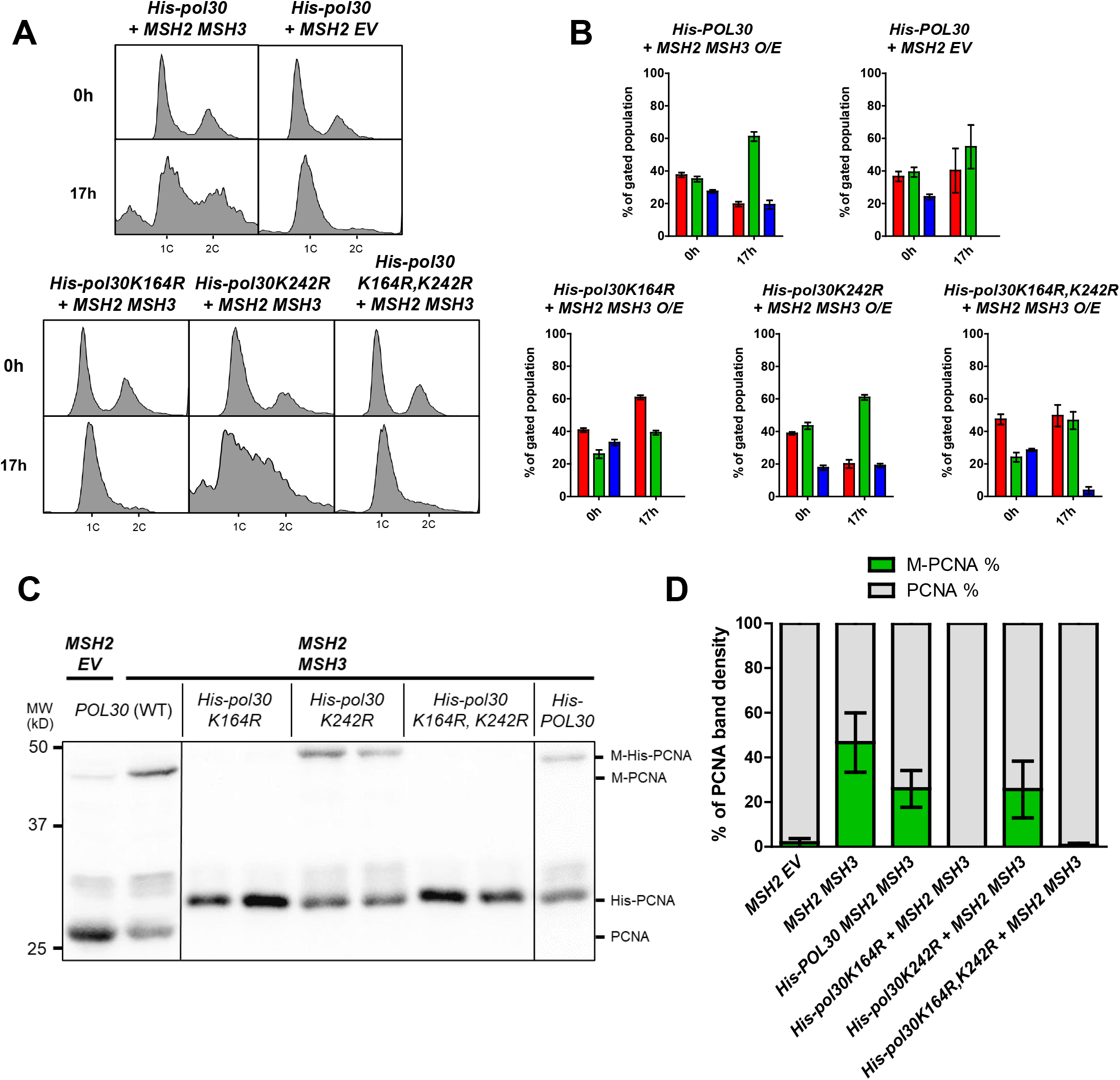
PCNA post-translational modification in the context of *MSH2 MSH3* overexpression occurs at the K164 residue. *MSH2* and *MSH3* or *MSH2* and empty vector were overexpressed following galactose induction in a *his-POL30, his-pol30K164R, his-pol30K242R,* or *his-pol30K164R+K242R,* background. Aliquots were collected and fixed for flow cytometry analysis before (0h) and after (17h) induction. After induction, cells were harvested for TCA extraction and anti-PCNA western blot. **(A)** Histograms are shown of chromosomal content of asynchronous populations before and after induction with galactose. Flow cytometry experiments were repeated at least three times, with at least two independent transformants. 1C indicates 1x DNA content; 2C indicates 2x DNA content. **(B)** Quantification of relative proportion of cells in different phases of the cell cycle. The percentage of cells in G1 (1C), G2/M (2C) or S (between 1C and 2C) phases was determined using BD FlowJo software (see Figure S1 for details). Plotted values correspond to data collected from at least three independent experiments. Error bars represent SEM. **(C)** PCNA Western Blot. Note that the PCNA and M-PCNA of *his-POL30* strains migrates at a higher weight due to the His-tag. These data are from a single blot, but not adjacent lanes, as indicated by the vertical lines. **(D)** PCNA Western Blot Quantification as described above.

The following observations suggest that this modification of PCNA is SUMO. First, the *MSH2 MSH3* overexpression-dependent PCNA modification exhibited the same mobility as the PCNA modification induced by high MMS levels (**Figure S5A**), which are known to promote PCNA sumoylation (102,103). Second, the modified band was not detected when TCA-precipitated His-PCNA was analyzed by western blot, using an α-ubiquitin (α-Ub) antibody (**Figure S5B**). Third, mass spectrometry of modified His-PCNA following *MSH2 MSH3* overexpression failed to detect ubiquitination (data not shown). Finally, the PCNA modification was abrogated in the absence of *SIZ1* (**Figure 8A,B**), which encodes the SUMO E3 ligase responsible for K164 sumoylation (104). Notably, the *MSH2 MSH3* overexpression-dependent cell cycle phenotype was also dependent on *SIZ1*. There was no S phase accumulation following *MSH2 MSH3* overexpression in *siz1Δ* (**Figure 8C,D, Figure S6**).

**Figure 8.**
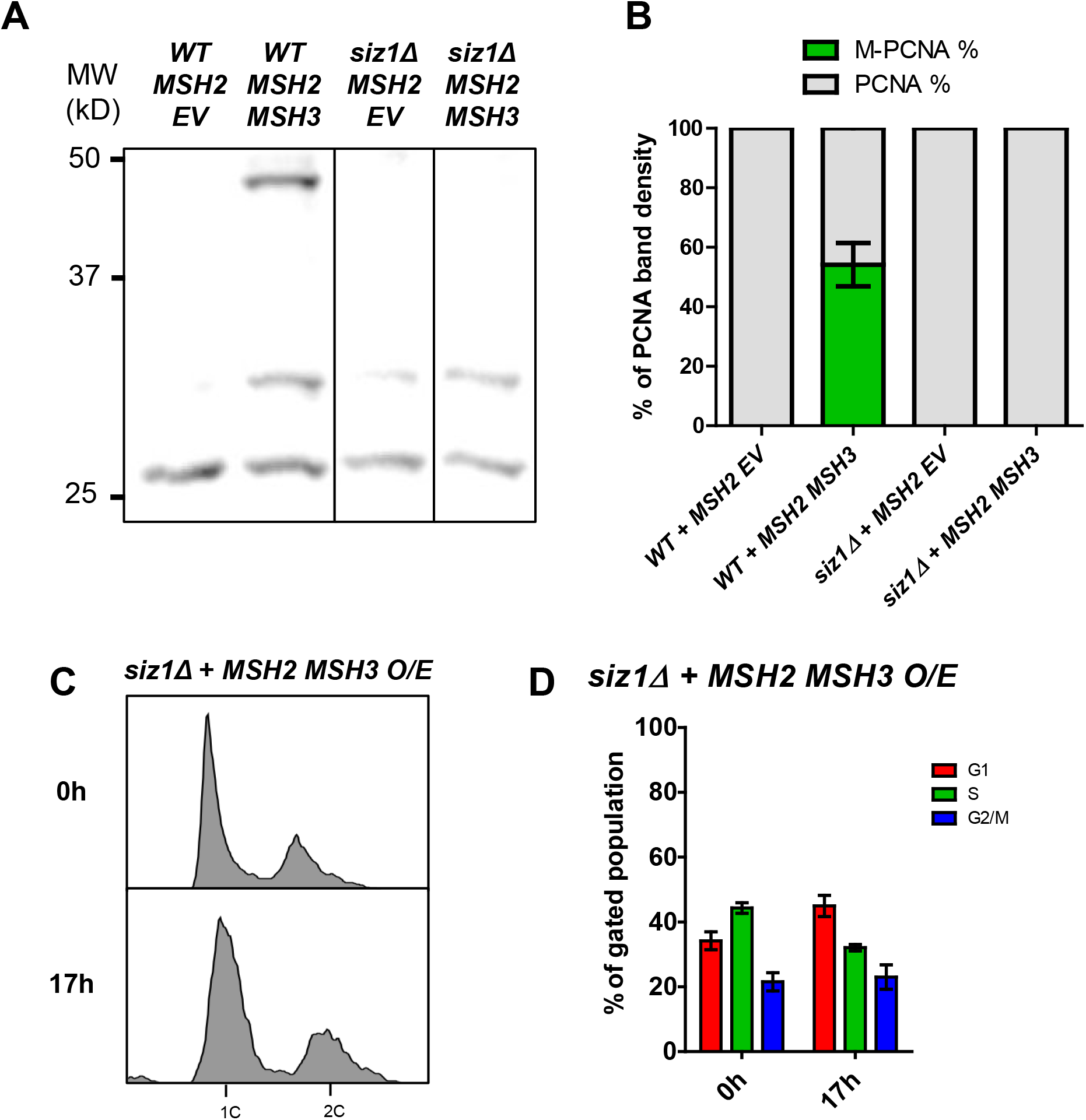
*MSH2 MSH3* overexpression-induced cell cycle defects are dependent on Siz1. *MSH2* and *MSH3* were overexpressed in a *siz1*Δ background, as previously described. Samples were collected for flow cytometry analysis before and after induction, and cells were harvested for Western Blot after induction. **(A)** Western Blot for PCNA. Modified PCNA is marked as M-PCNA. These data are from a single blot, but not adjacent lanes, as indicated by the vertical lines. **(B)** PCNA Western Blot quantification as previously described. **(C)** Flow cytometry analysis before and after induction. **(D)** Quantification of cell cycle analysis based on flow cytometry data.

The cell cycle analysis of these strains indicated that K164 is also required for the *MSH2 MSH3* overexpression-dependent accumulation of cells in S phase, while K242 was not essential. We note that the His-PCNA strain had a distinct cell cycle profile, with fewer cells in G2/M. Nonetheless, the accumulation of cells in the S phase remained clearly observable following *MSH2 MSH3* overexpression (**Figures 7A,B**).

### The *MSH2 MSH3* overexpression phenoype is *RAD9*-dependent

*MSH2 MSH3* overexpression leads to cell cycle delays, from which the cells recover, and post-translational modification of PCNA, all suggestive of activation of a cell cycle checkpoint response. Rad9, a cell cycle checkpoint protein involved in DNA damage signaling, is activated in response to impaired Okazaki fragment processing (87,105). We, therefore, overexpressed *MSH2 MSH3* in a *rad9Δ*. Deletion of *RAD9* was sufficient to suppress the cell cycle disruption caused by *MSH2 MSH3* overexpression (**Figure 9A, B**). The proportion of PCNA modified following *MSH2* MSH3 overexpression was also reduced in a *rad9Δ* background (**Figure 9C, D**), although there was some residual modification, ~25% of what was observed in *RAD9* cells. These results indicated that a Rad9-mediated cell cycle checkpoint response becomes activated in the presence of excess Msh2-Msh3, leading to cell cycle arrest and PCNA modification, potentially as a result of Msh2-Msh3 interference with OFM.

**Figure 9.**
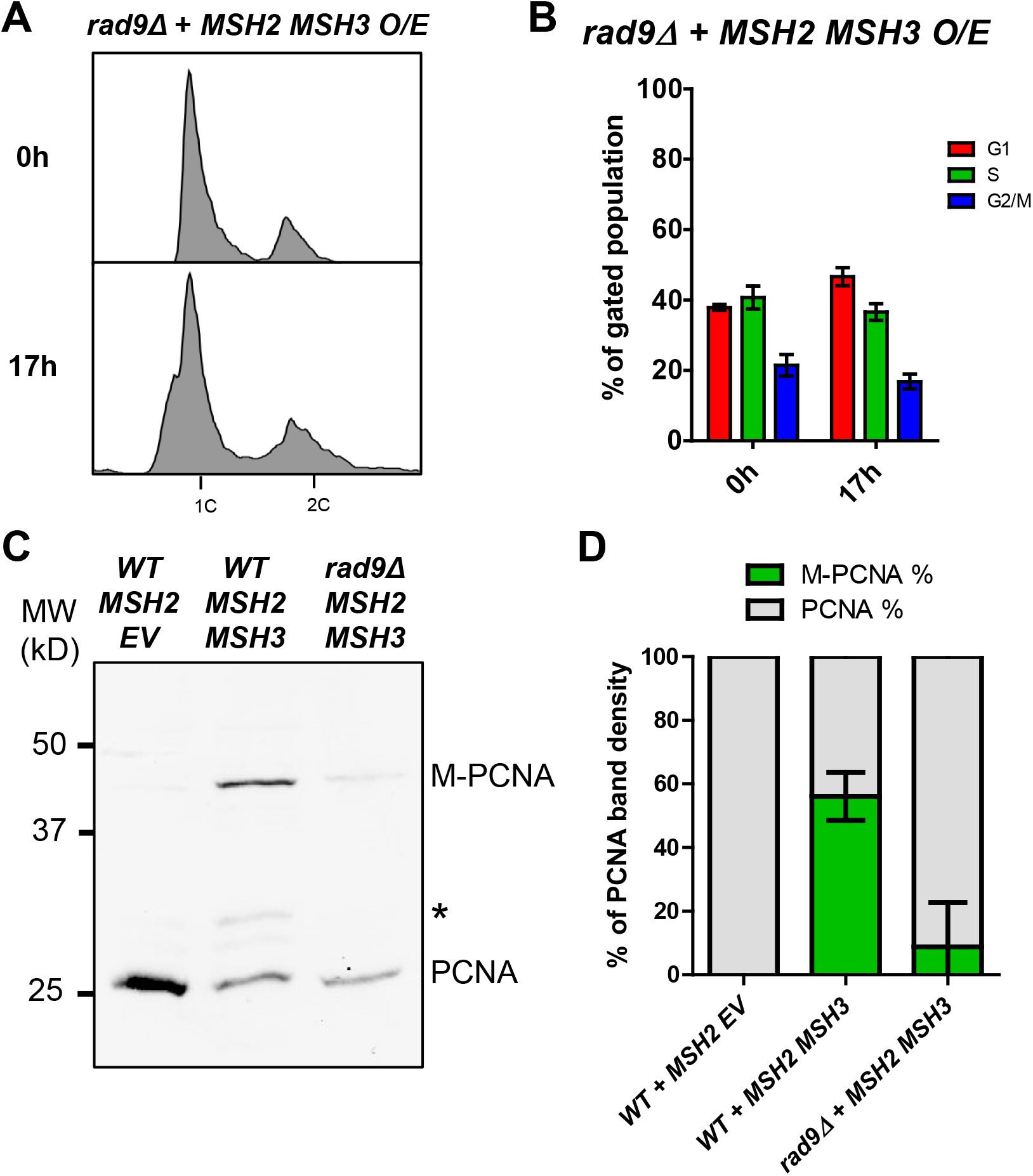
*MSH2 MSH3* overexpression-induced cell cycle defects are dependent on Rad9. *MSH2* and *MSH3* were overexpressed in a *rad9*Δ background, as previously described. Samples were collected for flow cytometry analysis before and after induction. **(A)** Flow cytometry analysis before and after induction. **(B)** Quantification of cell cycle analysis based on flow cytometry data. **(C)** Western Blot for PCNA. Modified PCNA is marked as M-PCNA. **(D)** PCNA Western Blot quantification as previously described.

### *ELG1* is required for PCNA modification and cell cycle defects caused by *MSH2 MSH3* overexpression

During lagging strand synthesis, PCNA must be loaded onto DNA at each Okazaki fragment by Replication Factor C (RFC), while PCNA unloading is carried out by a related complex in which Elg1 replaces the primary subunit in RFC, known as the Elg1-Replication Factor C-like Complex (Elg1-RLC) (106). Elg1 unloads both unmodified and SUMOylated PCNA but preferentially binds to SUMOylated PCNA (107). Elg1 unloading of PCNA at each Okazaki fragment is dependent on successful processing and ligation of the Okazaki fragment (108). Therefore, we decided to test the effects of *MSH2 MSH3* overexpression in an *elg1Δ* background. We found that the *MSH2 MSH3* overexpression-dependent cell cycle defect was abrogated in *elg1Δ* (**Figure 10, Figure S6**). We also observed reduced modification of PCNA (**Figure 10**). The requirement of *ELG1* for the cell cycle defect is consistent with the hypothesis that *MSH2 MSH3* overexpression interferes with Okazaki fragment processing *in vivo*.

**Figure 10.**
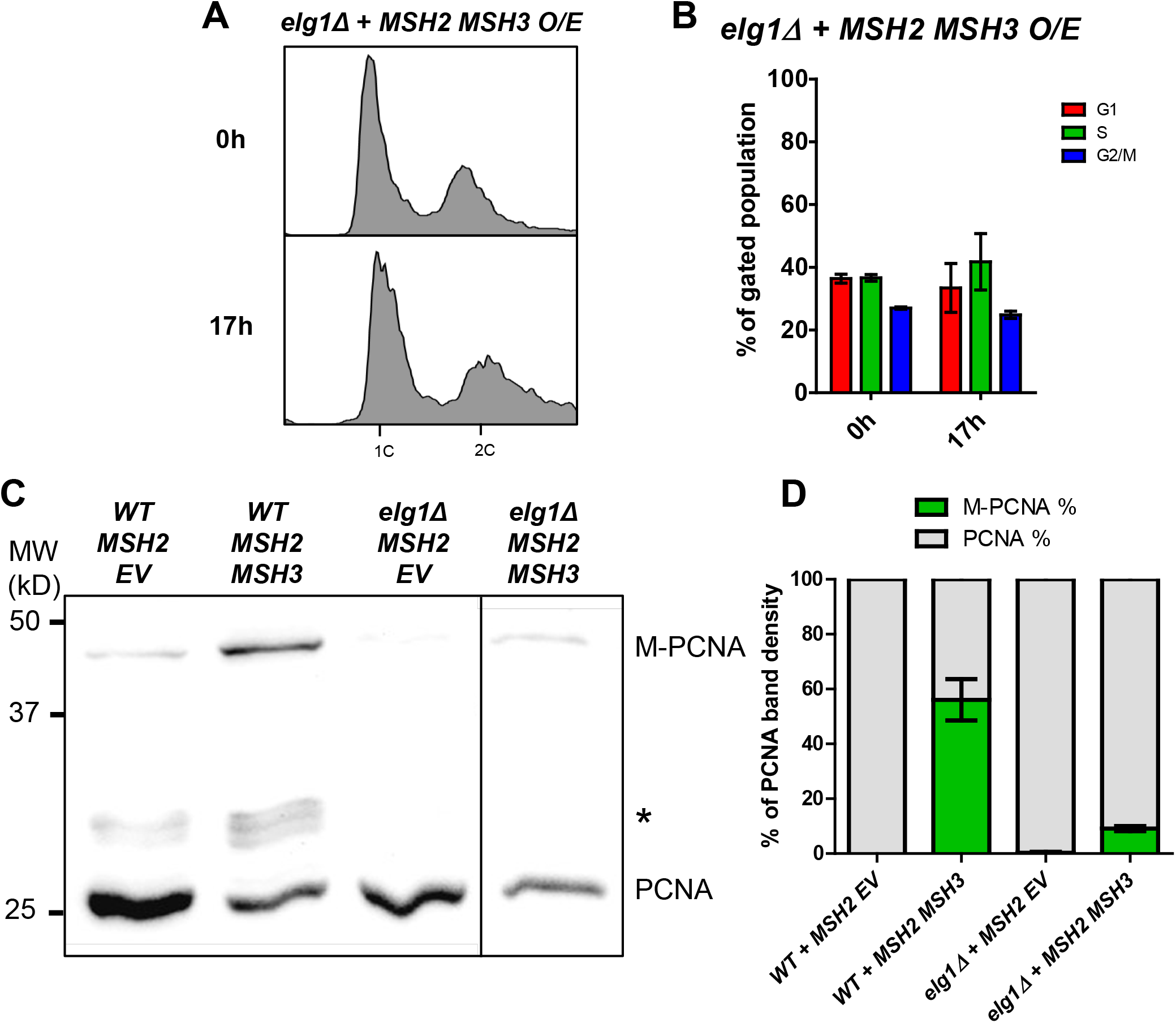
*MSH2 MSH3* overexpression-induced cell cycle defects are dependent on *ELG1*. *MSH2* and *MSH3* were overexpressed in an *elg1*Δ background, as previously described. Samples were collected for flow cytometry analysis before and after induction. **(A)** Flow cytometry analysis before and after induction. **(B)** Quantification of cell cycle analysis based on flow cytometry data. **(C)** Western Blot for PCNA. Modified PCNA is marked as M-PCNA. These data are from a single blot, but not adjacent lanes, as indicated by the vertical line. **(D)** PCNA Western Blot quantification as previously described.

### *msh3* alleles that disrupt ATP-binding and hydrolysis activities suppress disruption of cell cycle progression and PCNA post-translational modification

One possibility was that *in vivo*, excess Msh2-Msh3 binding to 5’ ssDNA flaps sterically interfered with Okazaki fragment maturation. Alternatively, Msh2-Msh3 bound to 5’ ssDNA flaps could promote aberrant Msh2-Msh3 activity that contributed to a signaling cascade leading to genome instability, similar to what is thought to occur in the presence of TNR structures (65,91). To distinguish between these possibilities, we tested the ability of ATPase-deficient *msh2* and *msh3* mutants to affect cell cycle progression. ATP is essential for the function and regulation of Msh2-Msh3 but is not required for DNA binding (13,14,63,64,66). Therefore, if Msh2-Msh3 simply binds to DNA structures and blocks Rad27^FEN1^, ATP binding, and/or hydrolysis should be dispensable for inducing the cell cycle defect. In fact, defects in these activities might even exacerbate the phenotype, as ATP hydrolysis contributes to Msh2-Msh3 turnover on the DNA (14,66). Alternatively, functional ATPase activity may be required to observe this effect, e.g., to recruit downstream proteins, as is the case for promoting TNR expansions (65,91).

To test these possibilities, we co-overexpressed *MSH2* and either *msh3G796A* or *msh3D870A* (64,91). These mutations disrupt the highly conserved Walker A motif (*msh3G796*), which mediates ATP binding, or the Walker B motif (*msh3D870*), which mediates ATP hydrolysis (64,68). Notably, the cell cycle defect was significantly less pronounced when either *msh3* allele was overexpressed compared to the overexpression of wild-type *MSH3* (**Figure 11A, C**). This suggests that full induction of the cell cycle defect requires Msh3 ATP binding and hydrolysis. Disruption of the Msh2 Walker A motif (*msh2G693D MSH3*) also suppressed the cell cycle defects when overexpressed (**Figure 11E**), indicating that ATP binding to both Msh2 and Msh3 is required for Msh2-Msh3 to disrupt cell cycle progression.

**Figure 11.**
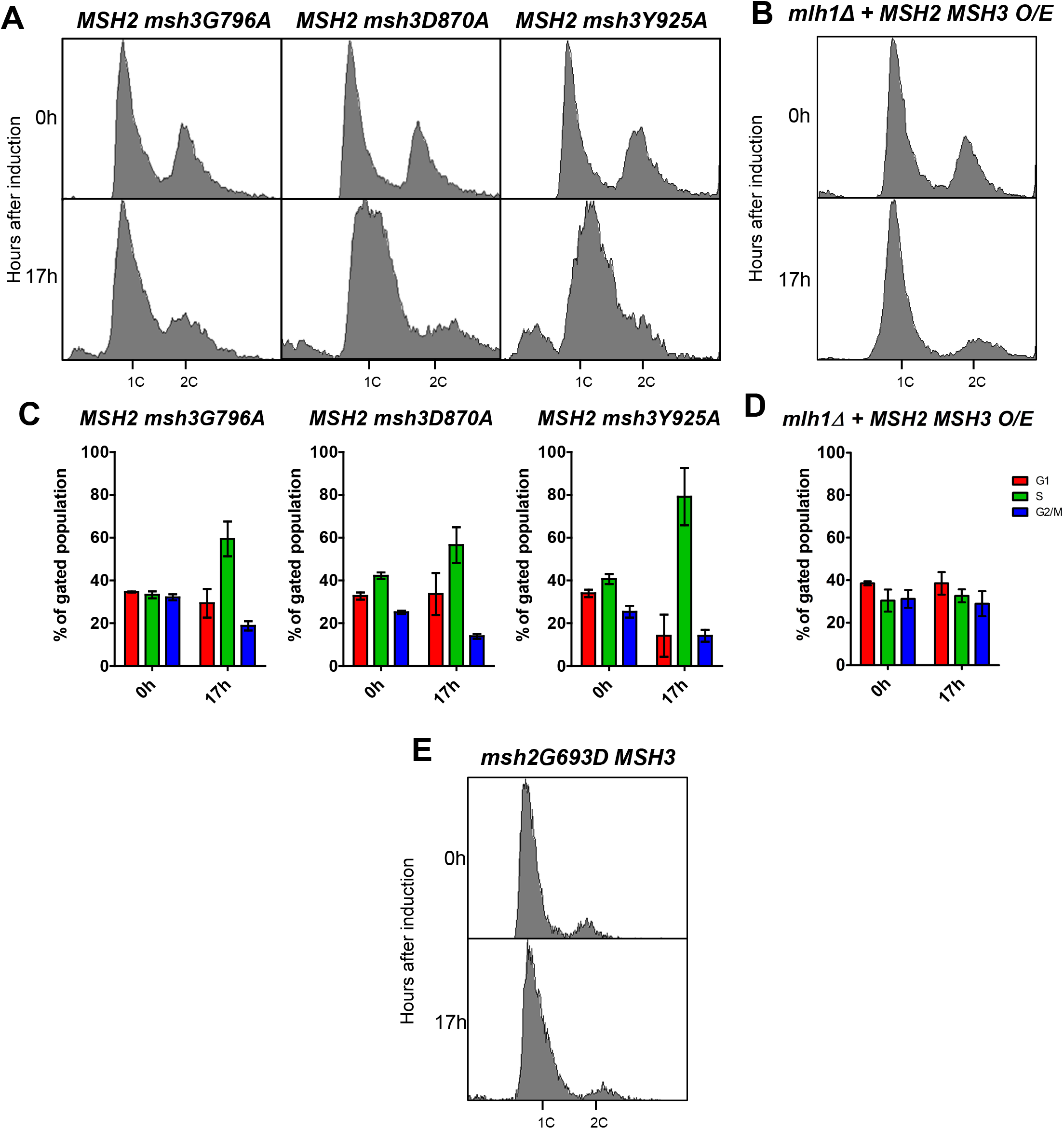
*MSH2 MSH3* overexpression-mediated cell cycle defect depends on MSH3 ATP binding and hydrolysis. *MSH2* (A-D) or *msh2G693D* (E) was co-overexpressed with *MSH3* (B, D) or *msh3* alleles **(A, C)** in a *msh3*Δ **(A, C)** or *mlh1*Δ **(B, D)** background. *MSH2* was constitutively overexpressed; for all others, expression was induced with galactose. Aliquots were collected at 0 and 17 hours after induction. Harvested cells were fixed, stained, and processed by flow cytometry. Histograms of the asynchronous population are shown.

We overexpressed *MSH2 msh3Y925A,* a separation-of-function allele that is defective in MMR but functional in 3’NHTR (64). Based on the human Msh2-Msh3 crystal structure, Y925 is predicted to regulate nucleotide occupancy of the nucleotide-binding pocket by pushing a conserved phenylalanine (F940) into the nucleotide-binding pocket (68). *In vitro,* Msh2-msh3Y925A retains ATP hydrolysis activity, but the kinetics of hydrolysis are significantly altered, indicating a defect in the regulation of ATP binding, ATP hydrolysis, and/or nucleotide turnover (**Figure S7**). When overexpressed, the *msh3Y925A* allele conferred cell cycle defects indistinguishable from the wild-type *MSH3* overexpression profile (**Figure A,C**). This is consistent with the hypothesis that ATP binding/hydrolysis by Msh2-Msh3 is required to impose cell cycle defects. However, regulation of this activity is less important, as observed for 3’ NHTR (64).

We also analyzed the effects of these *msh3* alleles on PCNA post-translational modification. When either the Walker A (*msh3G796A*) or Walker B (*msh3D870A*) motif was disrupted, post-translational modification of PCNA in response to *MSH2 msh3* co-overexpression was decreased to a similar extent (**Figure 12A,B**). Co-overexpression of *MSH2 msh3Y925A* promoted PCNA modification (**Figure S4**) at a level similar to wild-type, indicating that misregulated ATPase activity is sufficient for this effect. These results correlated with the intermediate effects of the Walker A and Walker B mutations and the wild-type effect of *MSH2 msh3Y925A* overexpression on the cell cycle progression phenotype (**Figure 11C**).

**Figure 12.**
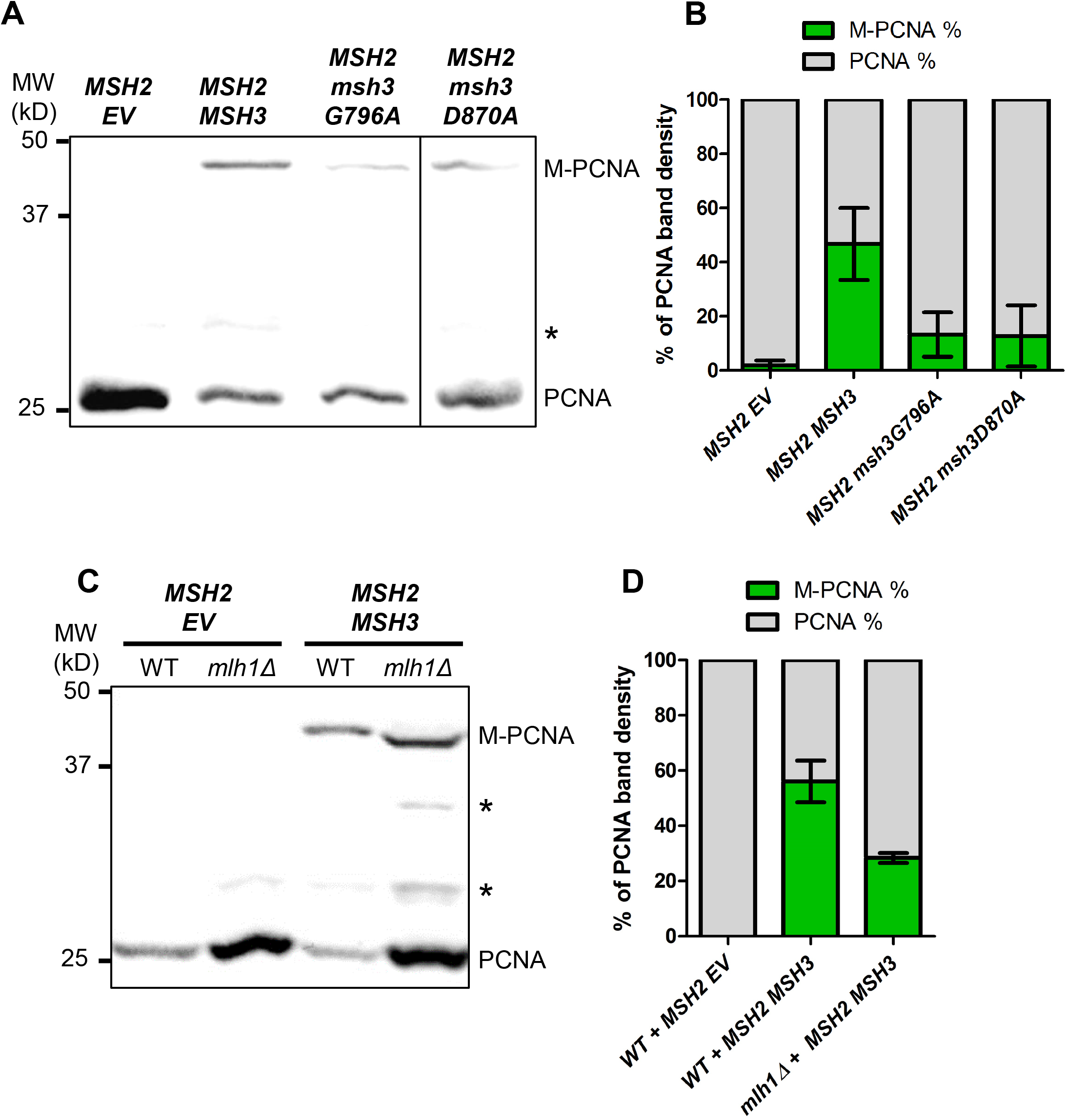
*MSH2 MSH3* overexpression induces PCNA post-translational modification in an ATP binding-and hydrolysis-dependent manner. *MSH2* and *MSH3* or *msh3* alleles were overexpressed in a *msh3*Δ **(A)** or *mlh1*Δ **(C)** background, as previously described. Western Blot for PCNA. Modified PCNA is marked as M-PCNA. The data in **(A)** are from a single blot, but not adjacent lanes, as indicated by the vertical line. **(B, D**) PCNA Western Blot quantification as previously described.

These data indicate that an active Msh2-Msh3-mediated pathway alters cell-cycle progression and induces post-translational modification of PCNA that might indicate replication stress and/or activation of a DNA damage response with Msh2-Msh3 levels are elevated.

### Downstream steps in MMR are required for cell cycle progression defects when MSH3 is overexpressed

The observation that Msh2-Msh3 ATP binding and hydrolysis activities are required to observe defects in cell cycle progression indicated that downstream steps in a Msh2-Msh3-mediated pathway might also be required to observe this phenotype. In MMR, Msh2-Msh3 DNA-binding leads to the recruitment of Mlh complexes and activation of their latent endonuclease activity (20,21,109–113). We tested whether *MLH1* is required for the cell cycle defect when *MSH2* and *MSH3* are co-overexpressed. We created a *mlh1Δ* strain, effectively inhibiting any downstream MMR activity by eliminating all three Mlh complexes: Mlh1-Pms1, Mlh1-Mlh2, and Mlh1-Mlh3. The *MSH2 MSH3* overexpression-dependent cell cycle defect was eliminated in the *mlh1*Δ background (**Figure 11B,D; Figure S6**). These results indicate that one or more Mlh complexes contribute to Msh2-Msh3-mediated replication stress that disrupts the cell cycle. Notably, we still observed enhanced PCNA modification under these conditions (**Figure 12C,D**), indicating the elevated Msh2-Msh3 is sufficient for this effect.

### Msh2-Msh3 modulates DNA Polymerase δ synthesis activity in vitro

We previously demonstrated that Msh2-Msh3 interferes with Rad27^FEN1^ and Cdc9^LigI^ activity when allowed to bind the relevant DNA substrates (28). Given that Msh2-Msh3 binds to ss/dsDNA junctions, we considered the possibility that Msh2-Msh3 interacting with different DNA structures might also impact DNA polymerase activity. As Pol δ is likely recruited in both Okazaki fragment processing and LP-BER, we tested Msh2-Msh3’s effect on Pol δ activity *in vitro* in the presence of a simple primer-template (synthesis) DNA substrate or a strand displacement substrate (**Figure 13**). We compared Msh2-Msh3 substrate binding efficiency on both DNA structures and found Msh2-Msh3 binds efficiently to both, albeit with lower affinity on the strand displacement substrate compared to the synthesis substrate (compare lanes 3 and 6, **Figure 13A**). Next, we assessed the ability of Msh2-Msh3 to modulate Pol δ synthesis on the synthesis substrate. Titration of Msh2-Msh3 into a primer extension reaction inhibited synthesis by Pol δ (**Figure 13B**). We observed similar inhibition on the strand displacement substrate (observe substrate retention in lanes containing Msh2-Msh3). However, in contrast to synthesis inhibition, we also observed a stimulation in the strand displacement synthesis products on a small subset of substrates. (**Figure 13B**). These results supported the hypothesis that Msh2-Msh3 can modify DNA metabolism pathways in a DNA structure-dependent manner, with variable impacts on genome stability.

**Figure 13.**
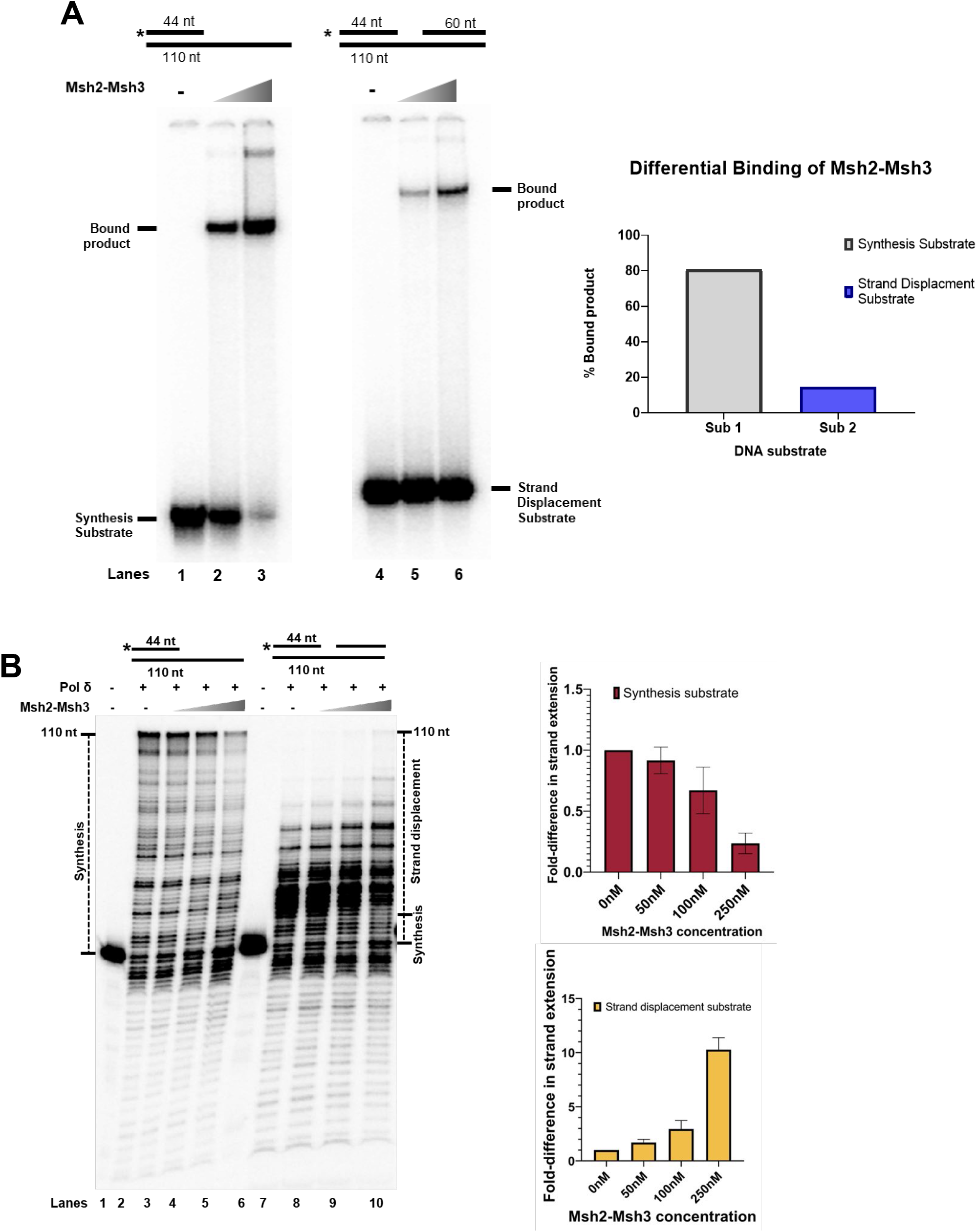
Msh2-Msh3 modulate DNA polymerase δ activity in a DNA substrate-dependent manner. **(A)** Msh2-Msh3 binds to the synthesis substrate (left) and the strand displacement substrate (right) with different affinities. Quantification is shown in right panel. **(B)** Msh2-Msh3 inhibits DNA polymerase δ in the presence of synthesis substrate (left) and stimulates its strand displacement synthesis activity (right).

## DISCUSSION

Msh2-Msh3 binds to a wide range of DNA structures (13,14,27,28,68,114,115). In this study, we demonstrated that Msh2-Msh3 could potentially interfere with *in vivo* DNA metabolism pathways that involve distinct DNA structures, including 5’ ssDNA flap intermediates. Even low levels of *MSH3* overexpression increased *in vivo* MMS sensitivity, and higher levels compromised progression through the S phase. This effect was not simply due to Msh2-Msh3 binding to DNA but rather required downstream steps, including ATPase activity and MLH complexes. *MSH3* overexpression appeared to trigger a *RAD9*-dependent DNA damage checkpoint response and modification of PCNA at K164.

### Maintaining the “right” amount of Msh2-Msh3 is important for genome stability

Altered Msh2-Msh3 expression, up or down, can be deleterious. Downregulation of Msh2-Msh3 is linked to tumorigenesis and cancer (38,47,116), while upregulation of Msh2-Msh3 promotes TNR expansions (91,92). At the same time, Msh complexes are differentially expressed among organisms and tissues (61,92,116–118). For instance, analysis of the abundance of different Msh complexes in actively proliferating murine tissues (including testis, spleen, and thymus) showed high expression of Msh6 but no Msh3; on the other hand, low proliferative tissues such as muscle, heart, and brain showed high *MSH3* expression levels (117). These differences suggest that proliferating cells produce enough Msh2-Msh3 to participate in processes that maintain genome stability (*e.g.,* MMR and 3’NHTR) but keep Msh3 at lower levels relative to Msh2 and Msh6 to limit aberrant DNA metabolic processes.

We previously demonstrated approximately equal levels of Msh2 and Msh6 levels in logarithmically growing cells, by quantitative immunoblotting (119). While Msh2 and Msh6 expression seem to depend on one another for stability, Msh3 is not stabilized by the presence of either Msh2 or Msh6 (120). Other studies measuring relative levels of Msh2, Msh3, and Msh6 by mass spectrometry (121) or quantitative immuno-purification (122) have observed an excess of Msh2 and Msh6 relative to Msh3. In this study, we assessed the transcript levels of untagged, endogenous *MSH2, MSH6,* and *MSH3* by qRT-PCR. Our results indicate that a wild-type yeast strain has a relative endogenous mRNA expression pattern of: Msh6,… Msh2 > Msh3. Overexpression of Msh3, which generates an imbalance of this distribution, was previously linked with strong mutator phenotypes in human cells (61,62). Consistent with this, when we overexpressed *MSH3* alone, we observed an increase in canavanine resistance, which primarily measures Msh2-Msh6 activity, indicating a decrease in Msh2-Msh6 function (**Table 1**). Decreased Msh2-Msh6 activity when Msh3 is overexpressed is presumably caused by a reduction in the formation of Msh2-Msh6 protein complexes, disrupting the “balance” between Msh2-Msh6 and Msh2-Msh3 complex formation. When we co-overexpressed *MSH2* and *MSH3* in budding yeast, we observed distinct phenotypes that included sensitivity to alkylating DNA damage (**Figure 3**), cell cycle delays (**Figures 4**, **5**), and an enhancement of post-translationally modified PCNA (**Figure 6**). Our results suggest that controlling the abundance of Msh3 is a mechanism by which cells can limit the interactions of Msh2-Msh3 with DNA structures, including 5’ ssDNA flaps, that modify Msh2-Msh3 function to promote genome instability. A similar correlation between Msh3 levels and TNR expansions was previously observed in a mouse model (92) and human cell lines (91). We predict that both the absolute levels of Msh2-Msh3 and the relative levels of Msh2-Msh3 versus Msh2-Msh6 are important to maintain the “right” balance of MMR activities in different cellular and genomic contexts.

### An MMR-like response is required for Msh2-Msh3-mediated cell cycle genomic instabilities

Initially, the *in vivo* effects of elevated *MSH3* or *MSH2 MSH3* expression suggested a simple model in which Msh2-Msh3 recognized and bound 5’ ssDNA flap structures as previously demonstrated *in vitro* (14,28), thereby blocking Rad27^FEN1^ activity *in vivo*. This would explain the MMS sensitivity, the defect in cell cycle progression through the S phase, and the modification of PCNA. However, two key observations, specifically the requirement for 1) Msh2-Msh3 ATPase activity and 2) the requirement for *MLH1*, indicated an active, Msh2-Msh3-mediated aberrant MMR-like response reminiscent of current models for Msh2-Msh3’s role in promoting TNR expansions (54,55,65,91,123).

Msh2-Msh3-mediated S phase accumulation and PCNA modification depended on Msh2-Msh3 ATP binding (Walker A) and hydrolysis (Walker B) activity. We and others have previously demonstrated that Msh2-Msh3 DNA binding does not require ATP (13,14,27,72). In fact, the presence of ATP promotes dissociation from DNA *in vitro* (14,27,72,115). Notably, the *msh2* and *msh3* Walker A mutations have dominant-negative effects *in vivo* (63,64); Msh2-msh3G693D inhibited ATP-dependent dissociation from DNA substrates *in vitro* (14), an observation interpreted to be a result of reduced Msh2-Msh3 turnover on the DNA. Therefore, DNA binding alone is not sufficient to produce the adverse effects of *MSH2 MSH3* overexpression. In contrast, *msh3Y925A*, predicted to alter the *regulation* of Msh2-Msh3 ATPase activity, did not reduce Msh2-Msh3-mediated interference with cell cycle progression. *msh3Y925A* also exhibited a dominant negative effect on MMR *in vivo* (64), indicating that this allele interfered with MMR but likely through a distinct mechanism. These observations are consistent with a model in which Msh2-Msh3 is not simply binding to 5’ ssDNA flaps and sterically hindering Rad27^FEN1^-mediated processing, although we predict that this capacity contributes to the cellular phenotypes.

Msh2-Msh3 ATPase activity is required for both Msh2-Msh3-mediated MMR and 3’ NHTR (64,66,124), although there are differential molecular requirements for the regulation of ATPase activity in these two pathways; *msh3Y925A* was defective in MMR but functional in 3’ NHTR (64). Further, Msh2-Msh3 nucleotide binding, hydrolysis, and turnover are differentially modulated by MMR versus 3’ NHTR DNA substrates. Msh2-Msh3 ATPase activity is similarly required to promote TNR expansions; mutations in the Msh2 Walker A or Msh3 Walker B motif disrupted TNR expansions in mammalian systems (65,91). Msh2-Msh3 binding to TNR DNA substrates altered nucleotide (ADP and ATP) binding and hydrolysis (27,72,91). We hypothesize that Msh2-Msh3 binding to non-canonical 5’ ssDNA flaps similarly alter Msh2-Msh3’s nucleotide binding/hydrolysis/turnover cycle and initiates an MMR-like response that disrupts OFM. Notably, Msh2-Msh3 may also be directly affecting DNA synthesis by Pol δ in OFM and/or LP-BER. We observed that Msh2-Msh3 either stimulated (synthesis substrate) or inhibited (strand displacement substrate) DNA Pol δ activity *in vitro* in a DNA structure-dependent manner. These observations, coupled with a differential affinity of Msh2-Msh3 for binding these substrates, predict a model in which Msh2-Msh3 binds and alters that conformation of the DNA structures to enhance or inhibit Pol δ activity. *In vivo*, Msh2-Msh3 could, when in sufficient quantities, similarly modulate Pol δ activity, altering and disrupting the kinetics of DNA synthesis.

Loss of Msh2-Msh3 ATPase activity also compromises its recruitment of Mlh complexes. Therefore, the requirement for ATP binding and, to a lesser extent, hydrolysis, may be related to the ability to recruit Mlh complexes. This is consistent with the requirement for *MLH1* to observe the *MSH2 MSH3* overexpression cell cycle phenotype. Loss of *MLH1* eliminates all three Mlh complexes. This suggests a model in which Msh2-Msh3 recruits one or more MLH complexes when bound to 5’ ssDNA flaps and that this interferes with Rad27^FEN1^-mediated pathways. Notably, all three Mlh complexes play a role in Msh2-Msh3-mediated TNR expansion (66,123,125–128). Based on our data, we suggest a model in which Msh2-Msh3 bound to a 5’ ssDNA flap intermediate recruits Mlh complexes, but Msh2-Msh3’s altered ATP binding/hydrolysis activity misregulates Mlh activation, similar to what has been proposed in TNR expansion studies, and interferes with Rad27-mediated OFM.

### A possible role of modified PCNA in Msh2-Msh3-mediated genome instability

The *MSH2 MSH3* overexpression-dependent modification of PCNA at K164 and the abrogation of this phenotype, as well as the cell cycle phenotype, in *elg1Δ* highlights a role for PCNA and its loading/unloading dynamics when the cells responds to elevated Msh2-Msh3 levels. PCNA plays a central role in DNA damage tolerance and repair signaling via post-translational modification. Monoubiquitination of PCNA at K164, catalyzed by the Rad6-Rad18 complex, recruits low-fidelity translesion synthesis polymerases (Pol η, Rev1, and Pol ζ) for potentially mutagenic lesion bypass (102,129–131). PCNA can be further poly-ubiquitinated at K164 by Ubc13-Mms2 and Rad5 to promote high-fidelity recombination (102,132,133). Alternatively, PCNA can be SUMOylated on residues K127 and/or K164, by the Ubc9-Siz1 complex, to prevent recombination through the recruitment of the anti-recombinase Srs2 (102–104,134–136). In this study, we showed that when excess Msh2-Msh3 is present *in vivo*, PCNA post-translational modification at K164 is enhanced (**Figure 6**, **7**). We were unable to detect PCNA ubiquitination by western blot (**Figure S5**) or by mass spectrometry (data not shown). However, the *siz1Δ* phenotypes and the requirement of *ELG1* for the cell cycle defect to be observed in the context of Msh2-Msh3 expression indicate that the modification is SUMO (**Figure 8,10; Figure S5**). SUMOylated PCNA is associated with the recruitment of Elg1(107).

The requirement for *ELG1*, which unloads unmodified and sumoylated PCNA to recycle PCNA during lagging strand synthesis (106–108,137), indicated that PCNA over-retention may be contributing to the *MSH2 MSH3* overexpression-dependent phenotype. Excess PCNA would inhibit OFM. We reason that if Msh2-Msh3 is interfering with cell cycle progression by 5’ flap processing, it might also be interrupting the normal cycling of PCNA. Such an interruption could result in the accumulation of SUMOylated PCNA at the replication fork, which would explain the enhancement of modified PCNA observed in **Figure 6**.

We also note that PCNA interacts with Msh complexes via their PIP-box motifs (138). The PCNA-Msh6 interaction recruits Msh2-Msh6 to the replication fork and plays a critical role in Exo1-independent MMR (19,119). Over-retention of PCNA on the DNA recruits increased Msh2-Msh6, resulting in elevated mutation rates and genomic instability (139,140). PCNA might similarly retain excess Msh2-Msh3 in *elg1Δ*. This could block Msh2-Msh3’s ability to recruit Mlh complexes; work in human Msh2-Msh3 demonstrated overlapping PCNA and MLH interaction motifs such that PCNA and Mlh complexes compete for binding to Msh2-Msh3 (141). As noted above, *MLH1* is required for the *MSH2 MSH3* overexpression phenotypes. (142).

### Model for MSH2 MSH3 overexpression-mediated genome instability

In this study, we demonstrated that overexpression of the Msh2-Msh3 complex in budding yeast could induce alkylation sensitivity (**Figure 3**), cell cycle progression delays (**Figures 4**, **5**), and a putative PCNA-mediated DNA damage response (**Figure 9**). These phenotypes required functional Msh2-Msh3 ATPase activity (**Figures 11**, **12**) and were abrogated in *rad9Δ*, *elg1Δ*, and *mlh1Δ* backgrounds (**Figures 9-12**). These data indicate that Msh2-Msh3 has the potential to disrupt many pathways in DNA metabolism, likely through its broad DNA binding capacity (**Figure 13**). Our findings further support the idea that Msh2-Msh3 binding alone is insufficient to determine between genome stability or instability outcomes. These features are similar to the requirements of Msh2-Msh3 in promoting TNR expansions (27,28,72,143,144), possibly suggesting a common mechanism for promoting genomic instability.

We propose the following model: (*i*) Increased Msh2-Msh3 abundance increases its ability to bind 5’ ssDNA flap intermediates during OFM or LP-BER, blocking efficient OFM and potentially activating a *RAD9*-dependent checkpoint response. (*ii*) Excess Msh2-Msh3 would also bind DNA substrates for Pol δ, either inhibiting (simple primer-template substrate) or enhancing (gapped substrate) its activity and altering the kinetics of DNA synthesis. (*iii*) Binding of a non-canonical DNA structure alters the ATP cycle within the Msh2-Msh3, potentially impacting its interaction with Mlh complexes. (*iv*) Msh2-Msh3 recruits Mlh complexes leading to aberrant activation of the Mlh via repeated rounds of nicking in an attempt to repair the DNA, leading to a checkpoint response. Alternatively, Msh2-Msh3 bound to 5’ ssDNA flap structures may misdirect Mlh endonuclease activity to the wrong strand, as demonstrated *in vitro* in the presence of TNR structures (145). It is clear that careful regulation of Msh2-Msh3 is critical for preventing aberrant or pathogenic outcomes in DNA metabolism, while retaining its advantageous genome stability functions.

## DATA AVAILABILITY

The data underlying this article will be shared on reasonable request to the corresponding author. All strains and plasmids will be made available.

## SUPPLEMENTARY DATA STATEMENT

Supplementary Data are available at NAR online.

## Supporting information

Medina-Rivera, Phelps et al Supplemental figures

Medina-Rivera, Phelps et al Supplemental tables

## ACKNOWLEDGEMENT

We are grateful for the support of the UB Confocal Microscope and Flow Cytometry Facility (CMFCF). We thank Katherine Casazza for technical assistance and for helpful discussions. We are grateful to Sarah Piacente and Jaime O’Connor for performing canavanine resistance assays and to Michaela (Cornaire) Radel for early MMS sensitivity assays.

