## Supplementary material for "Msh2-Msh3 interferes with DNA metabolism *in vivo*": Medina-Rivera, Phelps et al Supplemental figures

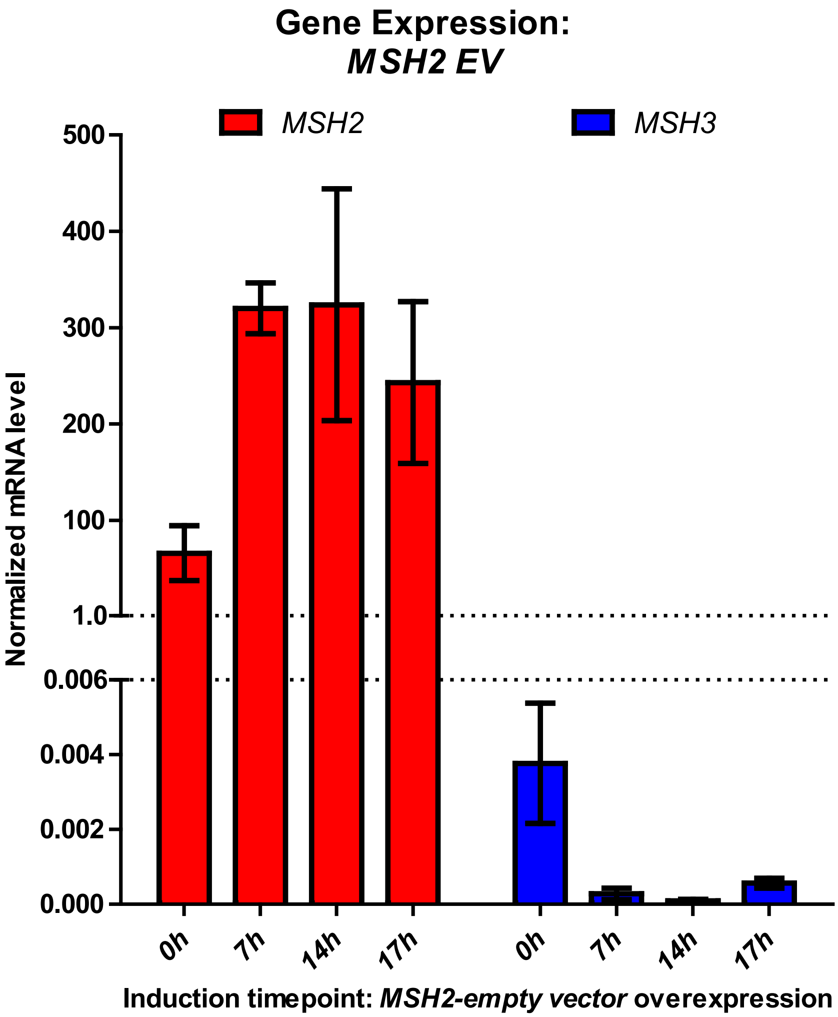


**Figure S1. *MSH2* + EV control shows no *msh3* mRNA.**

RNA was isolated from strains overexpressing *MSH2* under a constitutive promoter along with an empty vector plasmid (pJAS104). Endogenous mRNA levels of *MSH2* (pink, left) and *MSH3* (blue, right) were measured using qRT-PCR. All RNA levels were normalized to the reference gene *PDA1*. Bars represent mean ± SEM.

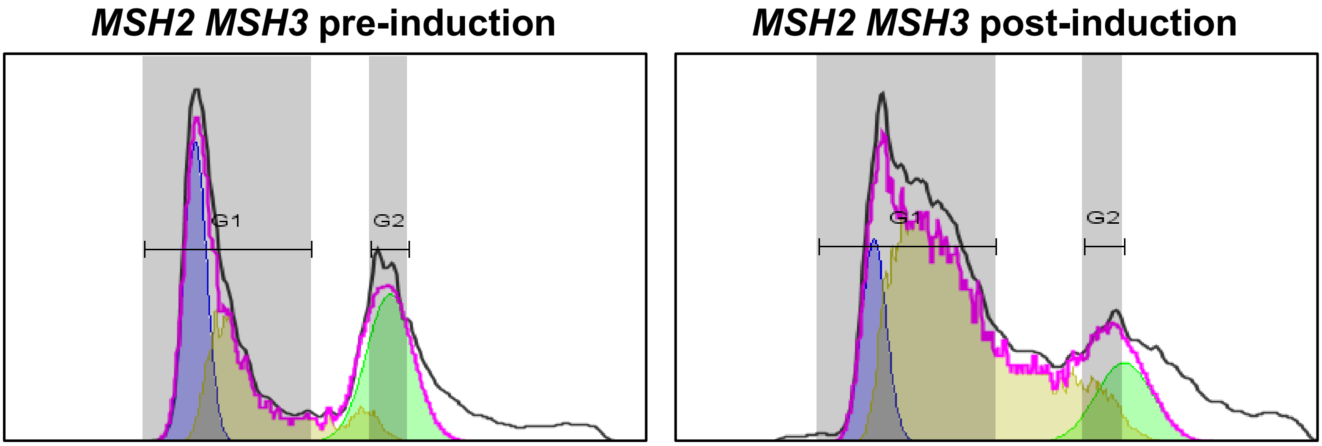


**Figure S2. Representation of BD FlowJo™ Quantification of Flow Cytometry data.**

Graphical representation of each time point accounts for a gated subset of the ten thousand total events recorded during each flow cytometry analysis. Gates P1, P2, and P3 excluded cellular debris and doublet events. Data shown on histogram in black represents cells in gate P3. Cell cycle analysis functionality inherent to the software was applied to P3 populations. Cells in G1, S, and G2/M are represented by areas under the blue (left), yellow (center), and green (right) curves, respectively. The pink line represents the sum of the curves. Ranges constraining the G1 and G2 peaks (vertical, gray) were tailored by sample.

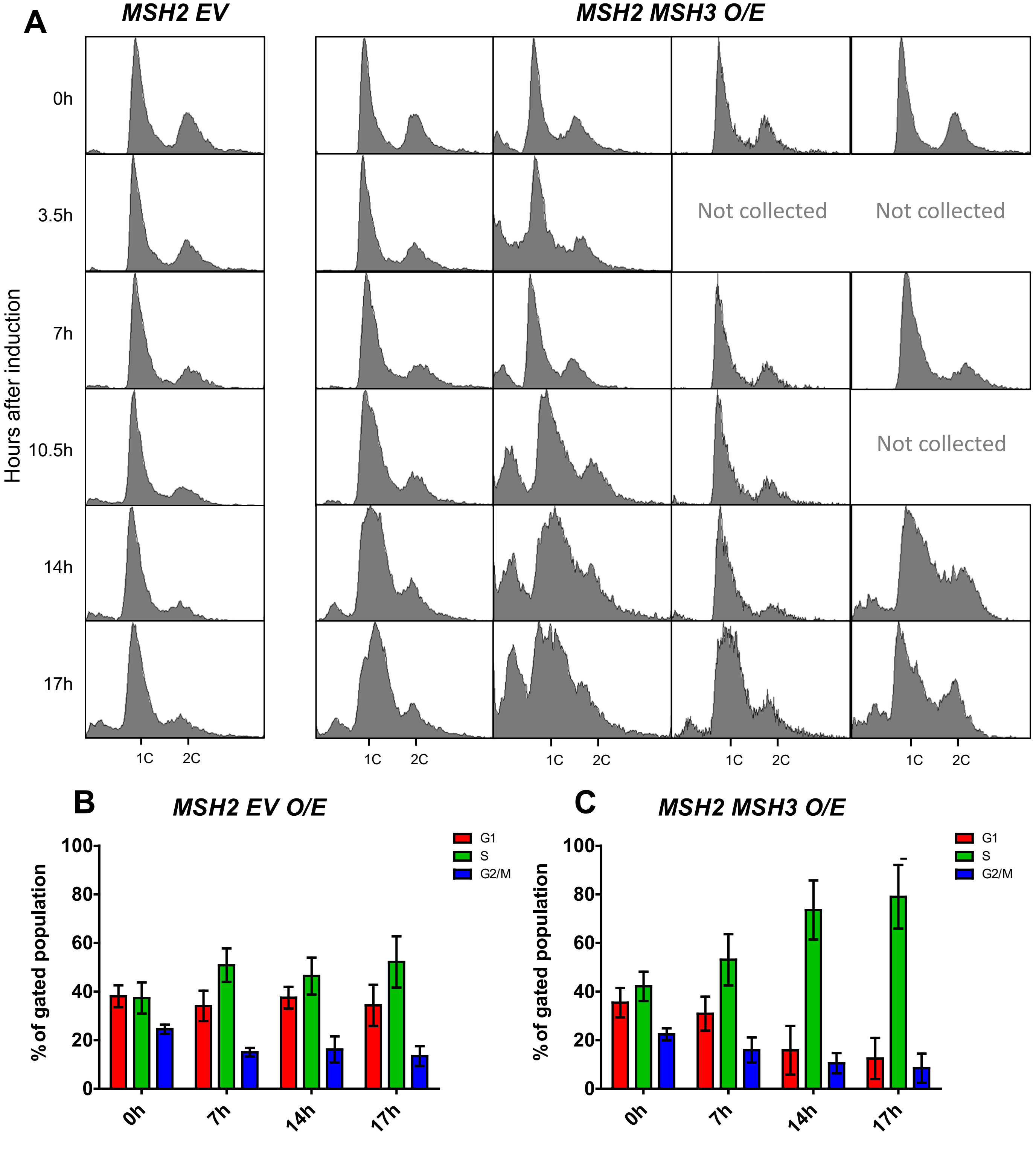


**Figure S3. *MSH2-MSH3* overexpression induces a delay in cell cycle progression.**
*MSH2* and *MSH3* were overexpressed following galactose induction in a *msh3Δ* background. Aliquots were collected at indicated hours after induction. (A) Histograms are shown of chromosomal content of asynchronous populations of *MSH2* + *empty vector (EV)* and *MSH2* + *MSH3* at several timepoints following addition of galactose. Flow cytometry experiments were repeated at least three times, with at least two independent transformants. 1C indicates 1x DNA content; 2C indicates 2x DNA content. (B and C) Quantification of relative proportion of cells in different phases of the cell cycle for *MSH2* + *EV* (B) or *MSH2* + *MSH3* (C). The percentage of cells in G1 (1C), G2/M (2C) or S (between 1C and 2C) phases was determined using FlowJo software (see Figure S1 for details). Plotted values correspond to data collected from at least three independent experiments. Error bars represent SEM.

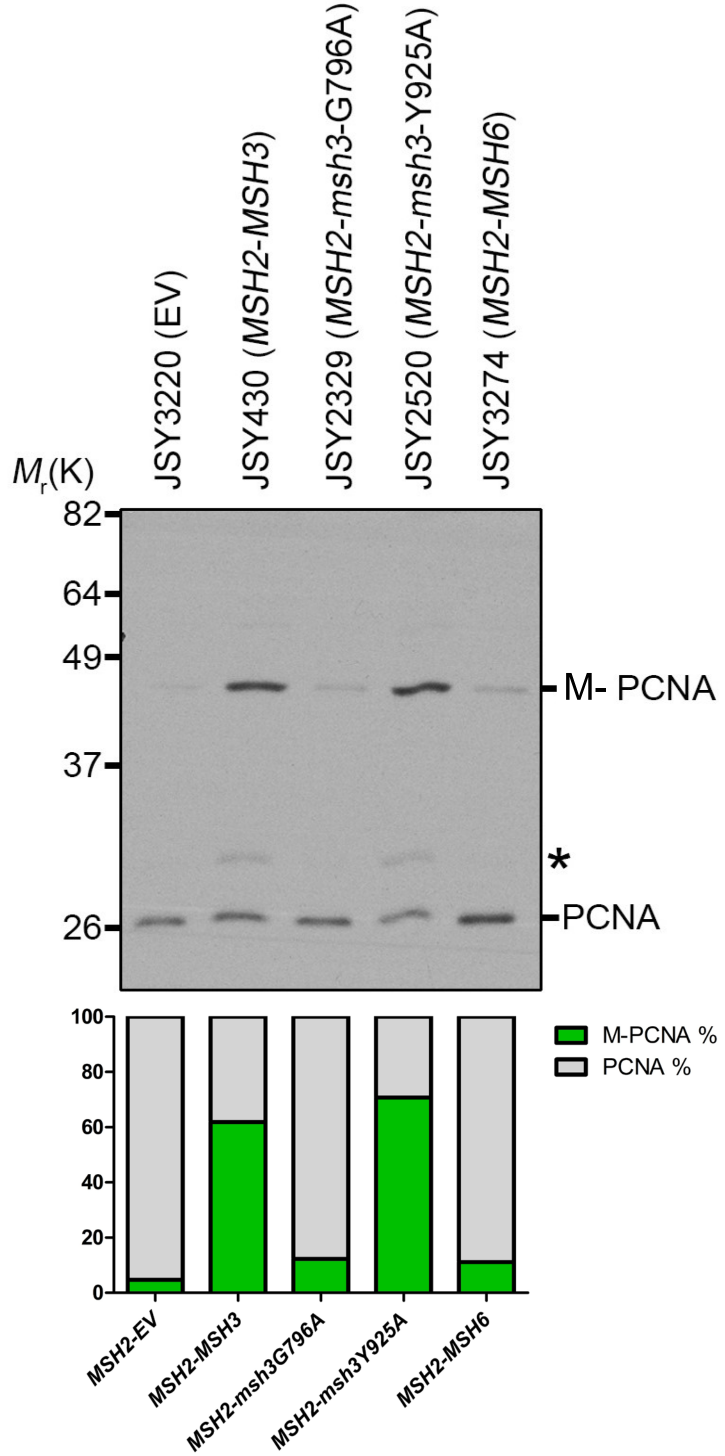


**Figure S4. ATP binding is required for *MSH2 MSH3* overexpression-induced PCNA modification, but proper regulation of ATP binding is not.**

*MSH2* and *MSH3* or *msh3* alleles were overexpressed in a *msh3Δ* background, as previously described. Samples were taken following induction and TCA protein extracts were prepared and the proteins were separated by 10% SDS-PAGE, transferred into a membrane and then probed with anti-PCNA. Modified PCNA is marked as M-PCNA. The asterisk indicates non-specific bands. Quantification of bands was done using ImageJ software and shown on the below graph as relative percentages of the total PCNA signal in that late on the blot shown (sum of modified & unmodified bands).

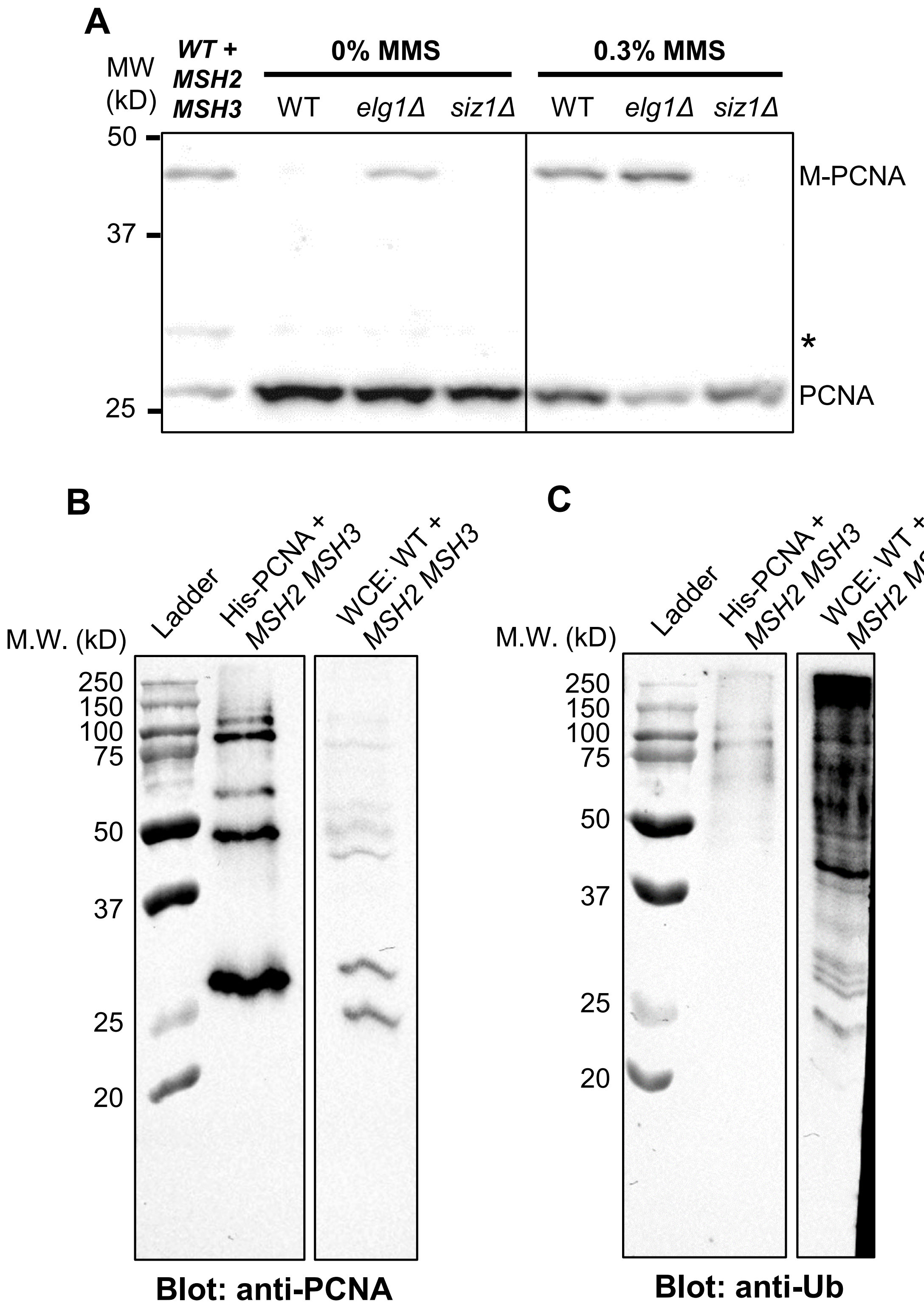


**Figure S5. Modified PCNA band may be SUMO-PCNA.**
**(A)** Wild-type, *elg1Δ*, and *siz1Δ* cells were treated with 0% or 0.3% MMS in mid-log phase for 90 minutes. Whole-cell extracts were collected by TCA precipitation and analyzed by Western blot for PCNA as described. **(B, C)** *MSH2* and *MSH3* were overexpressed as described in a strain carrying polyhistidine-tagged, wild-type PCNA *(His-POL30)*. Following induction, His-tagged PCNA was enriched using Ni-NTA agarose beads. Enriched His-PCNA samples ("His-PCNA + 2-3") were analyzed by Western blot. Following *MSH2 MSH3* induction in wild-type strain, whole-cell extracts (WCE) were collected by TCA precipitation as described and analyzed by Western blot ("WCE: WT 2-3"). **(B)** Samples were immunoblotted for PCNA. **(C)** Samples were immunoblotted for ubiquitin.

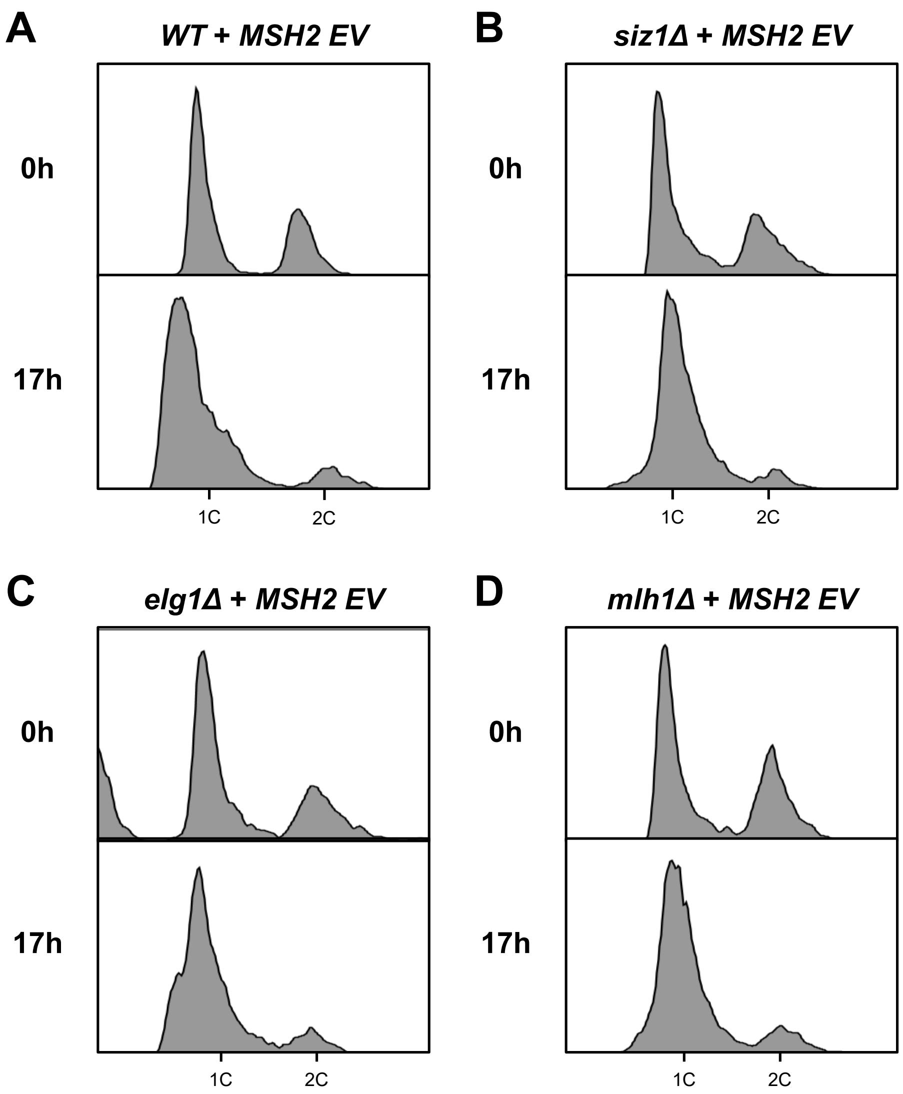


**Figure S6. Overexpression of *MSH2 EV* does not cause S phase accumulation in deletion strains.**

Representative hisotgrams of *MSH2* and empty vector (EV) overexpressed in S288C wild-type **(A)**, *siz1Δ* **(B)**, *elg1Δ* **(C)**, or *mlh1Δ* **(D)** backgrounds and induced as previously described. Cells were harvested for flow cytometry before and after induction.

A

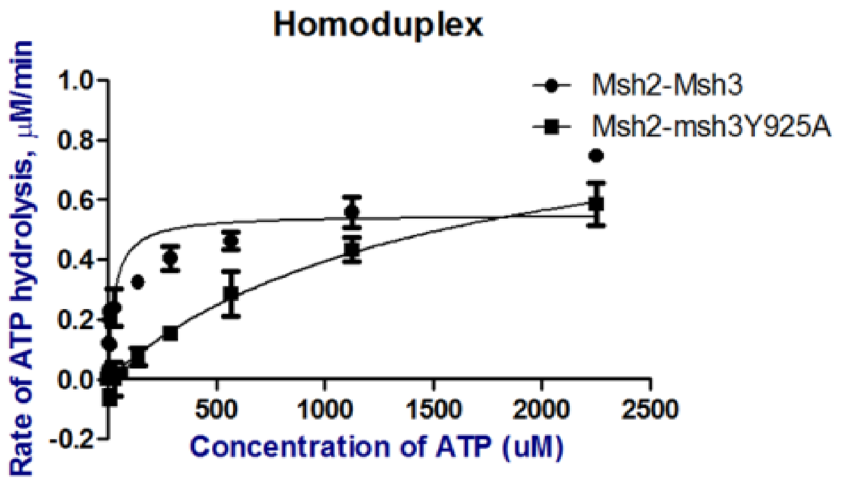


B

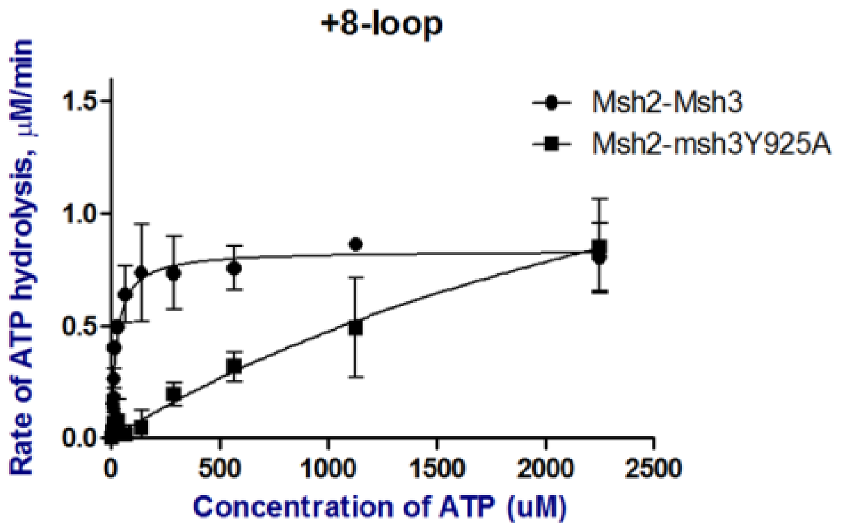


**Figure S7. ATPase activity of Msh2-Msh3 and Msh2-msh3Y925A in the presence of homoduplex and +8 loop (MMR) DNA substrates.**

The rate of hydrolysis was plotted against the concentration of ATP for wild-type Msh2-Msh3 (circles) and Msh2-msh3Y925A (squares). ATP hydrolysis was measured in the presenc of non-speciific homoduplex (**A**) or specific +8 loop (**B**) DNA substrates.
