## Supplementary material for "Msh2-Msh3 interferes with DNA metabolism *in vivo*": Medina-Rivera, Phelps et al Supplemental tables

### SUPPLEMENTARY TABLES

**Table S1 Plasmids used in this study**

| Plasmid | Allele | Origin; Marker | Reference |
|---|---|---|---|
| pMMR8 | *MSH2* | O/E; *TRP1* | (1) |
| pMMR20 | *MSH3* | O/E (Gal); *leu2D* | (1) |
| pJAS104 | Empty Vector | O/E (Gal); *leu2D* | (2) |
| pEAE218 | *MSH6* | O/E (Gal); *URA3* | (3) |
| pMME3 | *msh3D870A* | O/E (Gal); *leu2D* | This study |
| pCK94 or pMME2 | *msh3Y925A* | O/E (Gal); *leu2D* | (4), This study |
| pCK42 | *msh3G796A* | O/E (Gal); *leu2D* | (4) |
| pSP15 | *MSH3* | 2μ, *TRP1* | This study |
| pRS424 | Empty vector | 2μ; *TRP1* | (5) |
| pRS425 | Empty vector | 2μ; *LEU2* | (5) |
| pSP18 | *MSH3* | *ARS CEN; TRP1* | This study |
| pRS414 | Empty vector | *ARS CEN; TRP1* | (5) |

**Table S2 Strains used in this study**

| Strain Number | Genotype | Plasmid | Reference |
|---|---|---|---|

| | | | |
|---|---|---|---|
| FY23 (S288C background) | *ura3-52, leu2Δ1, trp1Δ63, his3Δ200, lys2Δ202, ura3Δ, leu2Δ, trp1Δ* | none | (6) |
| JSY905 | *msh3Δ* derivative of FY23 | none | This study |
| JSY3971-3972 | *msh3Y925A* derivative of FY23 | none | (4) |
| JSY3977-3979 | *msh3G796A* derivative of FY23 | none | (4) |
| JSY4418-4420 | FY23 *elg1Δ::KanMX* | none | This study |
| JSY4421-4424 | FY23 *mlh1Δ::KanMX* | none | This study |
| JSY4989-4991 | FY23 *rad9Δ::KanMX* | none | This study |
| JSY5077-5078 | FY23 *siz1Δ::KanMX* | none | This study |
| yb2062 (E133 derived) | *MATα, ade5-1, lys2-A12, trp1-289, his7-2, leu2-3,112, ura3-52, bar1* leu2*::His-POL30 (LEU2), pol30::TRP1* | none | (7) |
| yb2063 (E133 derived) | *MATα, ade5-1, lys2-A12, trp1-289, his7-2, leu2-3,112, ura3-52, bar1* leu2*::His-pol30K164R (LEU2), pol30::TRP1* | none | (7) |
| yb2064 (E133 derived) | *MATα, ade5-1, lys2-A12, trp1-289, his7-2, leu2-3,112, ura3-52, bar1* leu2*::His-pol30K242R (LEU2), pol30::TRP1* | none | (7) |
| yb2066 (E133 derived) | *MATα, ade5-1, lys2-A12, trp1-289, his7-2, leu2-3,112, ura3-52, bar1* leu2*::His-pol30K164R/K242R (LEU2), pol30::TRP1* | none | (7) |
| JSY4937-45 | *yb2062* leu2*::His-POL30 (leu2::hisG), pol30::trp1::hisG* | none | This study |
| JSY5007-12 | *yb2062* leu2*::His-pol30K164R (leu2::hisG), pol30::trp1::hisG* | none | This study |
| JSY5013-14 | *yb2062* leu2*::His-pol30K242R (leu2::hisG), pol30::trp1::hisG* | none | This study |
| JSY5015-27 | *yb2062* leu2*::His-pol30K164R/K242R (leu2::hisG), pol30::trp1::hisG* | none | This study |

***MSH3-HC* or *MSH3-LC***

| Strain Number | Genotype | Plasmid | Reference |
|---|---|---|---|
| JSY1789-1791<br>JSY4488-4490 | FY23 + LC Empty Vector | pRS414 | This study |
| JSY4183-4185 | FY23 + *MSH3-LC* | pSP18 | This study |
| JSY1786-1788 | FY23 + HC Empty Vector | pRS424 | This study |

| | | | |
|---|---|---|---|
| JSY4180-4182 | FY23 + *MSH3-HC* | pSP15 | This study |
| JSY4491-4493 | JSY905 + LC Empty Vector | pRS414 | This study |
| JSY4174-4176 | JSY905 + *MSH3-LC* | pSP18 | This study |
| JSY4384-4386 | JSY905 + HC Empty Vector | pRS424 | This study |
| JSY4171-4173 | JSY905 + *MSH3-HC* | pSP15 | This study |

***MSH2 MSH3*** **overexpression**

| **Strain Number** | **Genotype** | **Plasmid** |
|---|---|---|
| JSY1 | *ura3-52, trp1, leu2Δ1, his3Δ200, pep4::HIS3, prb1D1.6R, can1, GAL* | |
| JSY 430 | JSY1 | pMMR8, pMMR20 |
| JSY4387-4389 | JSY1 | pMMR8, pMME3 |
| JSY1439-4141 | JSY1 | pMMR8, pMME2 |
| JSY4136-4138 | JSY1 | pMMR8, pCK42 |
| JSY4133-4135 | JSY1 | pMMR8, pMMR20 |
| JSY4130-4132 | JSY1 | pMMR8, pEAO32 |
| JSY 1505 | *msh3Δ* derivative of JSY1 | |
| JSY 2329-2331 | JSY1505 | pMMR8, pCK42 |
| JSY 2520-2522 | JSY1505 | pMMR8, pCK94 |
| JSY4402-4404 | JSY1505 | pMMR8, pCK42 |
| JSY4399-4401 | JSY1505 | pMMR8, pMMR20 |
| JSY4396-4398 | JSY1505 | pMMR8, pMME3 |
| JSY4166-4168 | JSY1505 | pMMR8, pMME2 |
| JSY4390-4392 | FY23 | pMMR8, pMME3 |
| JSY4151-4153 | FY23 | pMMR8, pMME2 |
| JSY4148-4150 | FY23 | pMMR8, pCK42 |
| JSY4145-4147 | FY23 | pMMR8, pMMR20 |
| JSY4142-4144 | FY23 | pMMR8, pRS425 |
| JSY4393-4395 | JSY905 | pMMR8, pMME3 |
| JSY4163-4165 | JSY905 | pMMR8, pMME2 |
| JSY4160-4162 | JSY905 | pMMR8, pCK42 |
| JSY4157-4159 | JSY905 | pMMR8, pMMR20 |
| JSY4153-4156 | JSY905 | pMMR8, pRS425 |
| JSY4444-4446 | JSY4418 | pMMR8, pMMR20 |
| JSY4992-4997 | JSY4989, 4990 | pMMR8, pMMR20 |

| | | |
|---|---|---|
| JSY4447-4449 | JSY4421 | pMMR8, pMMR20 |
| JSY5082-5085 | JSY5077 | pMMR8, pMMR20 |
| JSY5079-5081 | JSY4942 | pMMR8, pMMR20 |
| JSY5028-5033 | JSY5010, 5011 | pMMR8, pMMR20 |
| JSY5034-5039 | JSY5013, 5014 | pMMR8, pMMR20 |
| JSY5040-5045 | JSY5015 | pMMR8, pMMR20 |

**Table S3 Oligonucleotides used for qRT-PCR**

| **Oligo** | **Gene** | **Sequence (5'–3')** | **Product size** |
|---|---|---|---|
| SO 351 | *MSH2* | GACAAGCAACAATCGGCTCTGGTT | 172 bp |
| SO 352 | | TCCATGGGATGCAACTTGGGTCTA | |
| SO 355 | *MSH3* | TGCGTACTGTTCTTTCCCGGATGT | 177 bp |
| SO 356 | | CTTGATTTGCTGGCACCTGGATCA | |
| SO 317 | *MSH6* | TACCTTCTGGCACACCGTCAAAGA | 107 bp |
| SO 318 | | TGCCTGTCTTTCCTCCTTGTGGAT | |
| SO 319 | *PDA1* | AACGCCAACCATCACAATTGGTCC | 173 bp |
| SO 320 | | ACGACTCGAAGGAAGATTCAGGCA | |

**Table S4: Oligonucleotides used for *in vitro* Polymerase δ assays**

| **Primer** | **Length** | **Sequence** |
|---|---|---|
| T1 | 110 | **3'**CAGGTGGGCGCGGTGGAGGACGAAGTTACACGACCTAGGATGTTGTTC TGCTTAAGCCTATGCTCCGGTCACGGCTGCACGGTCGGATTAAAGTTAGG TGG**5'** |
| U1 | 44 | 5' GTCCACCCGCGCCACCTCCTGCTTCAATGTGCTGGATCCTA3' |
| D1 | 60 | 5'CAGGTGGGCGCGGTGGAGGCCGTCACGGCTGGCAGGTCGGATTTAAG TTAGGTGGG3' |
